# The impact of uncertainty in hERG binding mechanism on *in silico* predictions of drug-induced proarrhythmic risk

**DOI:** 10.1101/2023.03.14.532553

**Authors:** Chon Lok Lei, Dominic G. Whittaker, Gary R. Mirams

## Abstract

**Background and Purpose:** Drug-induced reduction of the rapid delayed rectifier potassium current carried by the human Ether-à-go-go-Related Gene (hERG) channel is associated with increased risk of arrhythmias. Recent updates to drug safety regulatory guidelines attempt to capture each drug’s hERG binding mechanism by combining *in vitro* assays with *in silico* simulations. In this study, we investigate the impact on *in silico* proarrhythmic risk predictions due to uncertainty in the hERG binding mechanism and physiological hERG current model.

**Experimental Approach:** Possible pharmacological binding models were designed for the hERG channel to account for known and postulated small molecule binding mechanisms. After selecting a subset of plausible binding models for each compound through calibration to available voltage-clamp electrophysiology data, we assessed their effects, and the effects of different physiological models, on proarrhythmic risk predictions.

**Key Results:** For some compounds, multiple binding mechanisms can explain the same data produced under the safety testing guidelines, which results in different inferred binding rates. This can result in substantial uncertainty in the predicted torsade risk, which often spans more than one risk category. By comparison, we found that the effect of a different hERG physiological current model on risk classification was subtle.

**Conclusion and Implications:** The approach developed in this study assesses the impact of uncertainty in hERG binding mechanisms on predictions of drug-induced proarrhythmic risk. For some compounds, these results imply the need for additional binding data to decrease uncertainty in safety-critical applications.

## 1 Introduction

Reduction of the rapid delayed rectifier potassium current, *I*_Kr_, can lengthen the action potential (AP) which is associated with increased risk of cardiac arrhythmias, including Torsade de Pointes (Curran et al., 1995; Heist and Ruskin, 2010), although this risk is strongly modulated by multi-channel block (Mirams et al., 2011). The human Ether-à-go-go-Related Gene (hERG) encodes the pore-forming alpha subunit of the ion channel K_V_11.1 that conducts *I*_Kr_ (Sanguinetti et al., 1995). The hERG channel is highly susceptible to blockage or functional inhibition by a variety of pharmaceutical small molecules with a unique binding site (Sanguinetti and Tristani-Firouzi, 2006; Vandenberg et al., 2012), and its *in vitro* studies are part of the regulatory guidelines for proarrhythmic risk assessment (ICH, 2005). Recently, the Comprehensive in vitro Proarrhythmia Assay (CiPA) initiative and updates of regulatory guidelines consider a more nuanced characterisation of the proarrhythmic potential of a drug, by combining *in vitro* assays with *in silico* simulations that attempt to capture the details of each drug’s hERG binding mechanism, particularly the degree to which each drug is trapped (cannot unbind) when the channel closes (Sager et al., 2014; Li et al., 2019; ICH, 2022).

The inner cavity of the hERG channel is its principal drug binding site (Mitcheson, 2008). The molecular structure of the channel suggests that the channels can be blocked by pharmaceutical small molecules—we simply refer to these as “compounds” throughout—only when the channels are not closed (Butler et al., 2020), consistent with the observation that most of the compounds do not bind at negative (resting/repolarised) membrane potentials, when hERG channels are in a non-conducting closed state (Li et al., 2017). After binding to the channel, compounds unbind when channels are in different states and/or at different voltages (Mitcheson et al., 2000; Thouta et al., 2018). However, the unbinding process for some compounds can be impeded when the channels close, and the compounds remain bound, or “ *trapped* “ within the central cavity (Mitcheson et al., 2000; Stork et al., 2007; Windisch et al., 2011; Thouta et al., 2018), such as bepridil (Pareja et al., 2013) and dofetilide (Milnes et al., 2010); as opposed to, for example, cisapride (Milnes et al., 2010) and verapamil (Zhang et al., 1999) which unbind when the chancels close. Furthermore, according to the modulated receptor hypothesis, the difference in affinity determines the preferential binding of a compound to one of the states (Hille, 1977; Hondeghem and Katzung, 1977; Hondeghem, 1987; Carmeliet and Mubagwa, 1998), and its special case, the guarded receptor hypothesis, suggests the possibility compounds bind to a particular state only (Starmer and Courtney, 1986; Starmer et al., 1990, 1991). Both of these have been applied to models of hERG binding (Thurner et al., 2014; Lee et al., 2017; Veroli et al., 2013; Gomis-Tena et al., 2020).

To capture all the possible consequences of state-dependent binding within *in silico* models, all the above possibilities for drug binding to hERG should be considered. However, this raises the question of whether one size fits all—does the hERG binding model used in CiPA (Li et al., 2017) account for all the binding properties of interest, and do the model parameters reflect the underlying binding mechanisms? Furthermore, how sensitive is the proarrhythmic risk classification metric—the *torsade metric score* (Li et al., 2019)—to any uncertainty in the physiological hERG model and the binding mechanisms?

## 2 Methods

In this study, we designed a set of possible pharmacological binding models for the hERG channel that account for the known or postulated binding mechanisms (which we will describe in § 2.1–2.2). These models were calibrated to voltage-clamp electrophysiology data under the Milnes et al. (2010) protocol (§ 2.3.1) from Li et al. (2017, 2019), and plausible binding models that can explain the data well (or are approximately as good a fit to the data as the Li et al. (2017) model) were selected (§ 2.3.2). We then used these binding models with different hERG physiological models to calculate the “CiPA v1.0” risk metrics to assess the possible impact of different hERG binding mechanisms on such predictions (§ 2.4).

### 2.1 hERG physiological models

The basic physiological models of hERG describe the drug-free, control behaviour of the rapid delayed rectifier potassium current (*I*_Kr_) at physiological temperature and conditions. Two physiological models are used in this study, with the transition rates following *p*_*i*_ exp(*p*_*j*_*V*), where *p*_*i*_ and *p*_*j*_ are physiological model parameters taken from the literature. The first one, physiological model A, is shown in Figure 1A and is a six-state Markov model with two inactivated closed (IC) states, two closed (C) states, an inactivated (I) state, an open (O) state, and transition rates *a*_1_ to *a*_7_ and *b*_1_ to *b*_7_ (Li et al., 2017; Dutta et al., 2017). The second model, physiological model B, is a symmetric four-state model (Figure 1B) with transition rates *k*_1_ to *k*_4_, which is equivalent to a Hodgkin & Huxley-style model with one activation gate and one inactivation gate (the 37 °C average model in Lei et al., 2019a,b).

**Figure 1:**
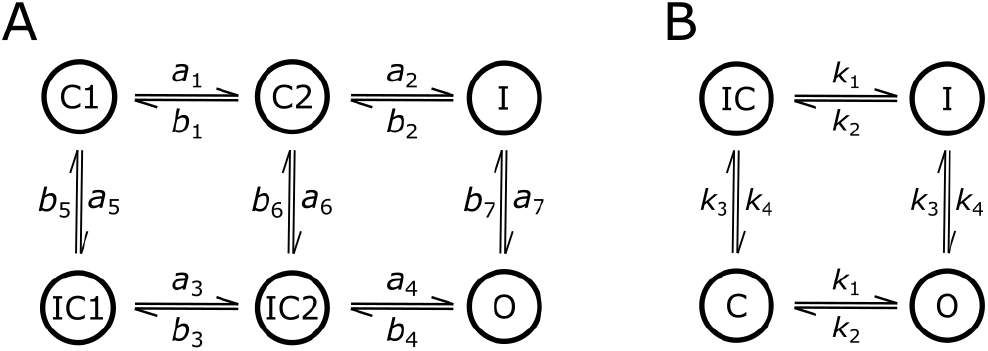
The base physiological models of the hERG channel considered in this study under drug-free conditions. **(A)** A six-state model from Li et al. (2017); Dutta et al. (2017) with the states renamed to match their respective physical states. **(B)** A symmetric four-state model from Beattie et al. (2018); Lei et al. (2019a). The models are used as the hERG model for studying drug effects.

### 2.2 Pharmacological binding models for hERG

To account for various known or proposed mechanisms for compounds binding to the hERG channel, a set of pharmacological binding models was designed as shown in Figure 2. The hERG physiological model is indicated in black (states I and O, and dots), representing either of the models from Figure 1.

**Figure 2:**
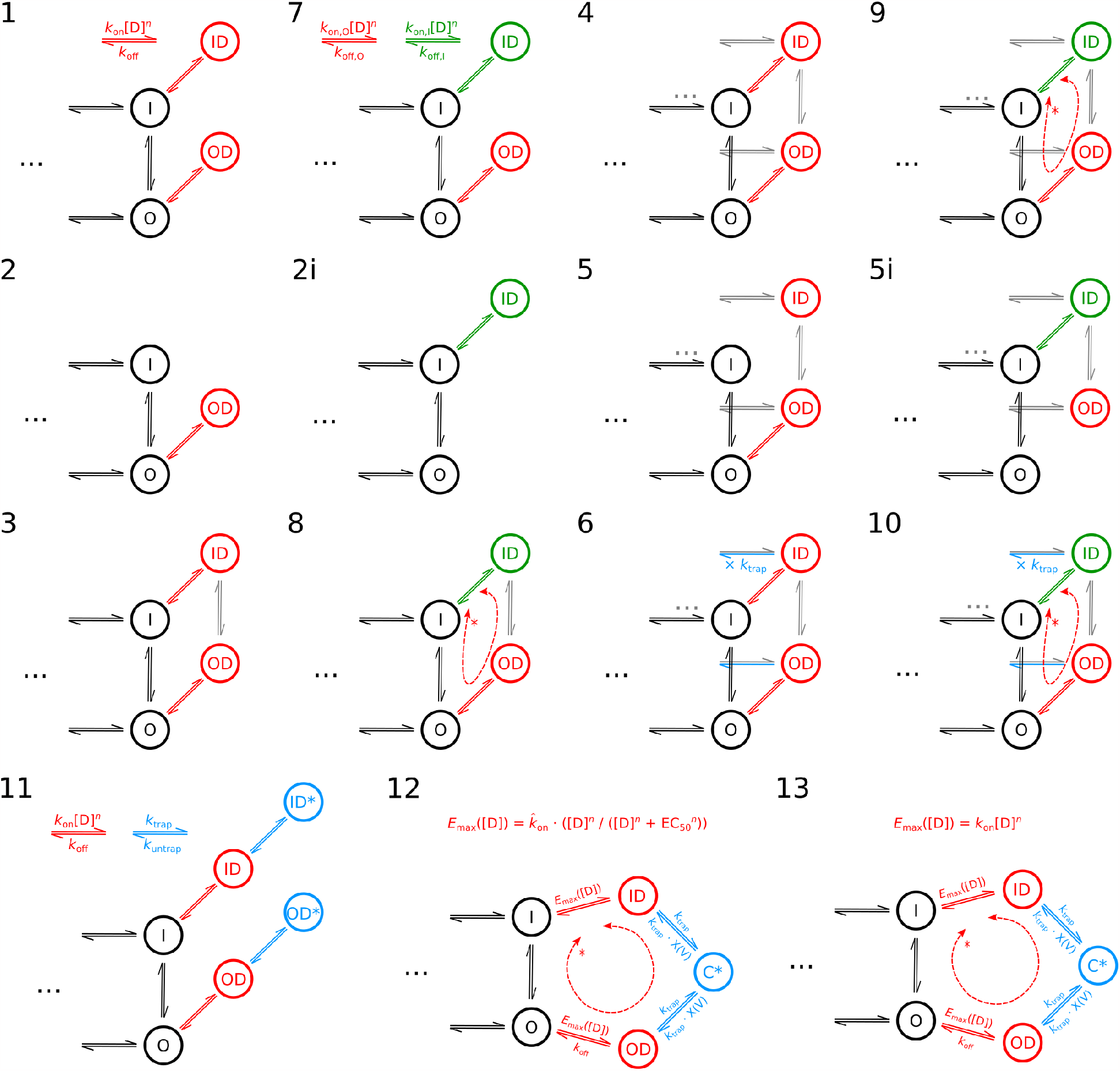
A set of pharmacological models representing different mechanisms of drug binding, where black (states I and O, and dots) is the physiological model of the hERG channel in Figure 1. Dashed double arrows indicate the rate marked with an asterisk is set by microscopic reversibility. Model 12 is identical to the pharmacological binding component in the CiPA v1.0 model (Li et al., 2017).

In Figure 2, Model 1 represents a compound that can bind to both open (O) and inactivated (I) states without being trapped in the binding pocket, and assumes the open and inactivated states share the same binding rate and unbinding rate, as shown in red. The binding rate is assumed to be proportional to the drug concentration [*D*] raised to the power of the Hill coefficient, *n*, and the association rate constant is *k*_on_; the unbinding rate is assumed to be a constant *k*_off_. Model 2 allows binding only to the open state; the inactivated version of Model 2 (Model 2i) allows binding only in the inactivated state, using the guarded receptor hypothesis. We have not observed any evidence that existing drugs bind at the resting potential (≈ 80 mV) when the channel is closed, therefore we have excluded binding models that bind only to the closed state. Model 3 is a variant of Model 1 where transition between the compound-bound states are also allowed, and happen at the same rate as the (unbound) O ⇌ I transitions.

Models 4 to 5i are the trapped equivalents of Models 1 to 2i, where the trapping component is indicated in grey, a ‘mirror image’ of the hERG physiological model with the same transition rates to admit the possibility of channels closing and preventing unbinding from CD or ICD states (not shown in the schematics). Model 6 relaxes the mirror trapping component of Model 4 by allowing an extra degree of freedom with a trapping rate factor *k*_trap_ multiplying the original transition rate. Models 7 to 10 allow an extra degree of freedom compared to Models 1, 3, 4, and 6 by assuming independent binding and unbinding rates for open and inactivated states—the modulated receptor hypothesis— whilst enforcing microscopic reversibility by specifying rates indicated by an asterisk as a function of the other rates in the closed loop (Colquhoun et al., 2004). Model 11 introduces independent trapped states for the open and inactivated compound-bound states with transition rates *k*_trap_ and *k*_untrap_, which is the ‘intermediate encounter complex’ model in Windley et al. (2016).

Finally, Model 12 is the same model (equations) as the CiPA v1.0 model proposed by Li et al. (2017)—the *reference model* of the study but we will refit parameters in this study. In Model 12, rather than the drug binding rate being linearly proportional to [*D*]^*n*^, instead it saturates, and this is represented using a Hill equation:

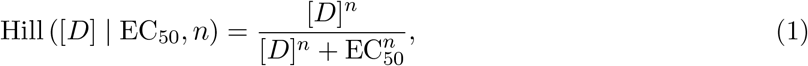

where EC_50_ is a half maximal effective concentration; in this case, when [*D*] = EC_50_, the binding rate is half of its maximum rate. Note that the binding rate parameter for Model 12 has a unit of ms^*−*1^, so we denote it as 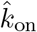 to differentiate it from *k*_on_ in the other binding models (which have units of ms^*−*1^ nM^*−n*^). When [*D*] ≪ EC_50_, Model 12 reduces to the usual linear binding rate, *k*_on_[*D*]^*n*^; we consider this case separately as Model 13. Furthermore, Models 12 and 13 assume the untrapping rate follows a sigmoid function of the membrane voltage *X*(*V*) = (1 + exp ((*V*_1*/*2,trap_ − *V*)/6.789))^*−*1^ instead of the voltage dependence of the control condition—a mirror image of the hERG physiological model; they also assume a fixed trapping rate *k*_trap_ = 3.5× 10^*−*5^ ms^*−*1^, with *V*_1*/*2,trap_ altering the degree of trapping (Li et al., 2017).

We also included two additional models: Models 0a and 0b, as basic and standard ‘conductance block’ models for drug effects. We consider an ‘all-state-blocker’ model that has binding and unbinding rates, *k*_on_[*D*]^*n*^ and *k*_off_, for all channel states. The degree of block is then independent of state occupancy, and is equivalent to having a modulating fraction of unavailable channels or ‘*b* gate’ such that we scale (multiply) the drug-free current by (1 − *b*). *b* itself follows the equation

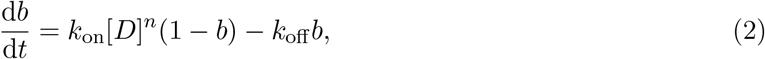

with an initial condition of *b* = 0 at the time a compound is first introduced, we call this Model 0b. Model 0a is a simple conductance scaling model which is equivalent to assuming Model 0b is an instantaneous process, i.e. the degree of block *b* is set immediately to the steady state of Eq. (2):

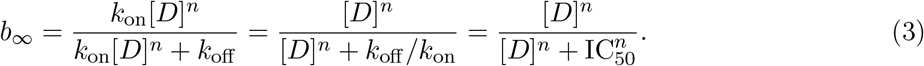

The ratio *k*_off_ */k*_on_ in Model 0b is known as the dissociation constant and is equivalent to IC 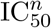 in the Hill equation in Model 0a. All the binding model parameters that need to be calibrated on a compound-specific basis are listed in Table 1.

**Table 1:** The model parameters for all the binding models in Figure 2. The asterisk indicates the rate parameter was determined by the microscopic reversibility.

| Model | Binding parameters |  |  |  |  |  | Trapping parameters |  |
| --- | --- | --- | --- | --- | --- | --- | --- | --- |
| 0a | $n$ | $IC_{50}$ | | | | | | |
| 0b, 1, 3, 4 | $n$ | $k_{on}$ | $k_{off}$ | | | | | |
| 2, 5 | $n$ | $k_{on,O}$ | $k_{off,O}$ | | | | | |
| 2i, 5i | $n$ | $k_{on,I}$ | $k_{off,I}$ | | | | | |
| 6 | $n$ | $k_{on}$ | $k_{off}$ | | | | $k_{trap}$ | |
| 7 | $n$ | $k_{on,O}$ | $k_{off,O}$ | $k_{on,I}$ | $k_{off,I}$ | | | |
| 8, 9, 10 | $n$ | $k_{on,O}$ | $k_{off,O}$ | $k_{on,I}$ | * | | | |
| 11 | $n$ | $k_{on}$ | $k_{off}$ | | | | $k_{trap}$ | $k_{untrap}$ |
| 12 | $n$ | $\hat{k}_{on}$ | $k_{off}$ | $EC_{50}$ | | | $V_{1/2,trap}$ | |
| 13 | $n$ | $k_{on}$ | $k_{off}$ | | | | $V_{1/2,trap}$ | |

### 2.3 Data and Statistical Analysis

#### 2.3.1 Electrophysiology data

The voltage-clamp electrophysiology data were taken for the 28 CiPA training and validation compounds (Li et al., 2017, 2019) where manual patch-clamp experiments were performed on HEK293 cells stably expressing hERG1a subunit at 37 °C. Data were collected using a modified Milnes’ protocol (Milnes et al., 2010); the protocol had 10 sweeps measured with 10 ms time point interval, and each repeat consisted of 1 s (with leak step) at the resting potential (−80 mV), followed by a long voltage step to 0 mV for 10 s, before returning to the resting potential for 14 s. For the details of the experiments, please refer to Li et al. (2017).

#### 2.3.2 Calibration of pharmacological binding models

We calibrated/fitted each of the pharmacological binding models in Figure 2 with each of the hERG physiological models in Figure 1 independently to the voltage-clamp electrophysiology data for each compound. The available data have been normalised to the control current, giving the *fraction of unblocked current* which reveals the change in current due to the drug binding. Furthermore, only the data at the voltage step 0 mV (time interval 1.1–11 s) for the 10 sweeps were used; the remaining data at the resting potential (−80 mV) will have almost no current, giving little information about the drug binding. All models, except Model 0a, were calibrated by minimising the root-mean-square difference (RMSD) of the percentage current between the model output and the mean experimental data from 10 sweeps for four different concentrations of a compound, giving 4 × 10 × 990 = 39600 data points for each compound. The optimisation was performed with logarithmic-transformed model parameters, except *V*_1*/*2,trap_ in Models 12 and 13. This allows a better search across a wide parameter range, especially for the rate parameters. The optimisation was performed using the covariance matrix adaptation-evolution strategy (CMA-ES) algorithm (Hansen, 2006) in PINTS (Clerx et al., 2019), and was repeated 10 times from different initial guesses sampled from wide boundaries (*k*_on_, *k*_off_ ∈ [10^*−*7^, 1] ms^*−*1^, *k*_trap_, *k*_untrap_ ∈ [10^*−*9^, 10^3^] ms^*−*1^, 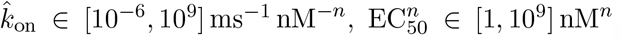, *V*_1*/*2,trap_ ∈ [0, 200] mV, and *n* ∈ [0.2, 2]) to ensure we arrived at a global minimum. The calibration was repeated for all 28 compounds listed in Li et al. (2019).

Model 0a is a special case of Model 0b where drug binding is approximated as instantaneously reaching steady state. Therefore Model 0a should not be calibrated to transient experimental data. Hence, the parameters of Model 0a were directly taken as the steady state of Model 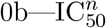 of Model 0a in Table 1 was calculated as the dissociation constant *k*_off_ */k*_on_ of Model 0b, as Eq. (3) suggests.

However, since each pharmacological binding model in Figure 2 was designed to model a specific mechanism, not all the models are expected to be able to describe all the compounds that were tested. Therefore each calibrated binding model was assessed by comparing its fitted RMSD to the RMSD of bootstrapped samples of the data which were computed as follows: since each of 10 repeated experiments at each concentration was independent, for one compound across all four concentrations, there are 10^4^ permutations. Therefore, we randomly selected (with replacement) one trace out of the ten available for each concentration to obtain a set of 4 traces for all concentrations; the RMSD of this set to the mean of the experimental data was computed, giving the RMSD of a “bootstrapped” data trace. This process was repeated 1000 times to get a range of RMSDs based on these bootstrapped samples of the data. This essentially compares how well the models fit to the mean data relative to an individual data trace’s fit to the mean data, providing a compound-dataset-specific measure of goodness-of-fit. A pharmacological binding model was classified as a *plausible model* if its RMSD was either (a) smaller than the maximum (excluding outliers) RMSD of the bootstrap samples of the data or (b) within 20 % of the RMSD of the CiPA v1.0 reference model.

### 2.4 AP models, *q*_net_ and torsade metric scores

To assess the impact of the choice of hERG binding model on predictions of drug-induced proarrhythmic risk, we adapted the approach used in Li et al. (2019). We used the optimised CiPA v1.0 AP model by Dutta et al. (2017) for predicting the effects on AP due to different hERG binding mechanisms. We replaced the “dynamic hERG-binding model” in the AP model with one of the successfully calibrated pharmacological binding models in Figure 2. To account for multiple ion channel block effects, the Hill equation characterised by the half-inhibition concentration (IC_50_) and the Hill coefficient (*n*)— Model 0a—was used to model the drug effects on three other currents, the fast sodium current (*I*_Na_), the late sodium current (*I*_NaL_), and the L-type calcium current (*I*_CaL_), using the reported median values from Li et al. (2019).

The hERG physiological model can be either of the models shown in Figure 1. However, to use the new Figure 1B hERG physiological model, we re-calibrated the *I*_Kr_ conductance, in control (drug-free) conditions, by matching the AP duration at 90 % repolarisation (APD_90_) at 0.5 Hz pacing (−80 A/F stimulus amplitude and 0.5 ms duration) after 1000 paces at (quasi-)steady state. We obtained a new *I*_Kr_ conductance of 0.0912 pA/pF for the hERG physiological model B in the AP model that resulted in the same APD_90_ as the CiPA v1.0 model.

The hERG-binding-model-replaced-AP models were used to calculate the metric *q*_net_ following the definition in Dutta et al. (2017). Pacing at 0.5 Hz (−80 A*/*F stimulus amplitude and 0.5 ms duration) was initialised from the steady state under control-conditions and continued for 1000 paces after compound addition, at multiples of each compound’s maximum therapeutic concentrations (*C*_max_). The *q*_net_ metric was defined as the net charge over one beat carried by *I*_Kr_, *I*_NaL_, *I*_CaL_, the transient outward potassium current (*I*_to_), the slow rectifier potassium current (*I*_Ks_), and the inwardly rectifying potassium current (*I*_K1_). The metric was computed by integrating the sum of the six currents between two consecutive stimuli (with time step 0.01 ms) using the trapezium rule. The proarrhythmic risk prediction was made using the *torsade metric score*, defined as the mean *q*_net_ value averaged at 1×, 2×, 3×, 4× *C*_max_ (Li et al., 2019).

Finally, following Li et al. (2019), an ordinal logistic regression model (all-threshold variant, Rennie and Srebro, 2005) was used to estimate the new drug-induced proarrhythmic risk thresholds for low-, intermediate-, and high-risk categories, using the torsade metric as the feature. The classifier was trained with L2 regularisation using only the training compounds and was solved with the L-BFGS-B algorithm in SciPy (Virtanen et al., 2020). The two thresholds for separating (a) the low-risk category from intermediate/high and (b) high from low/intermediate risks were calculated using

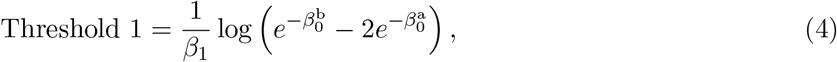

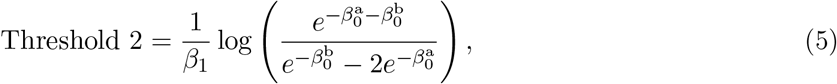

where 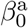 and 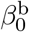 are the intercepts, and *β*_1_ the linear coefficient (of the torsade metric), of the linear equations that map the torsade metric through a logistic function to the cumulative probabilities of the risk categories within the logistic regression model.

## 3 Results

### 3.1 Multiple binding mechanisms can explain the same drug effects

All binding models (Figure 2) were calibrated to the experimental data for each compound independently and compared against the maximum RMSD for bootstrapped samples of the data. Figure 3 shows the calibrated binding models (left) and their RMSDs to the mean experimental data (right) across all binding models, for three example compounds: dofetilide (top), terfenadine (middle), and verapamil (bottom). The percentage current plots in Figure 3 (left) show *only* the effect of the drug over time for the 10 pulses of holding potential (0 mV) in CiPA-Milnes’ protocol. Regardless of the physiological models (A, squares and B, circles), the results of the calibration of the binding models were similar. The results of all the remaining compounds are shown in Supplementary Figures.

**Figure 3:**
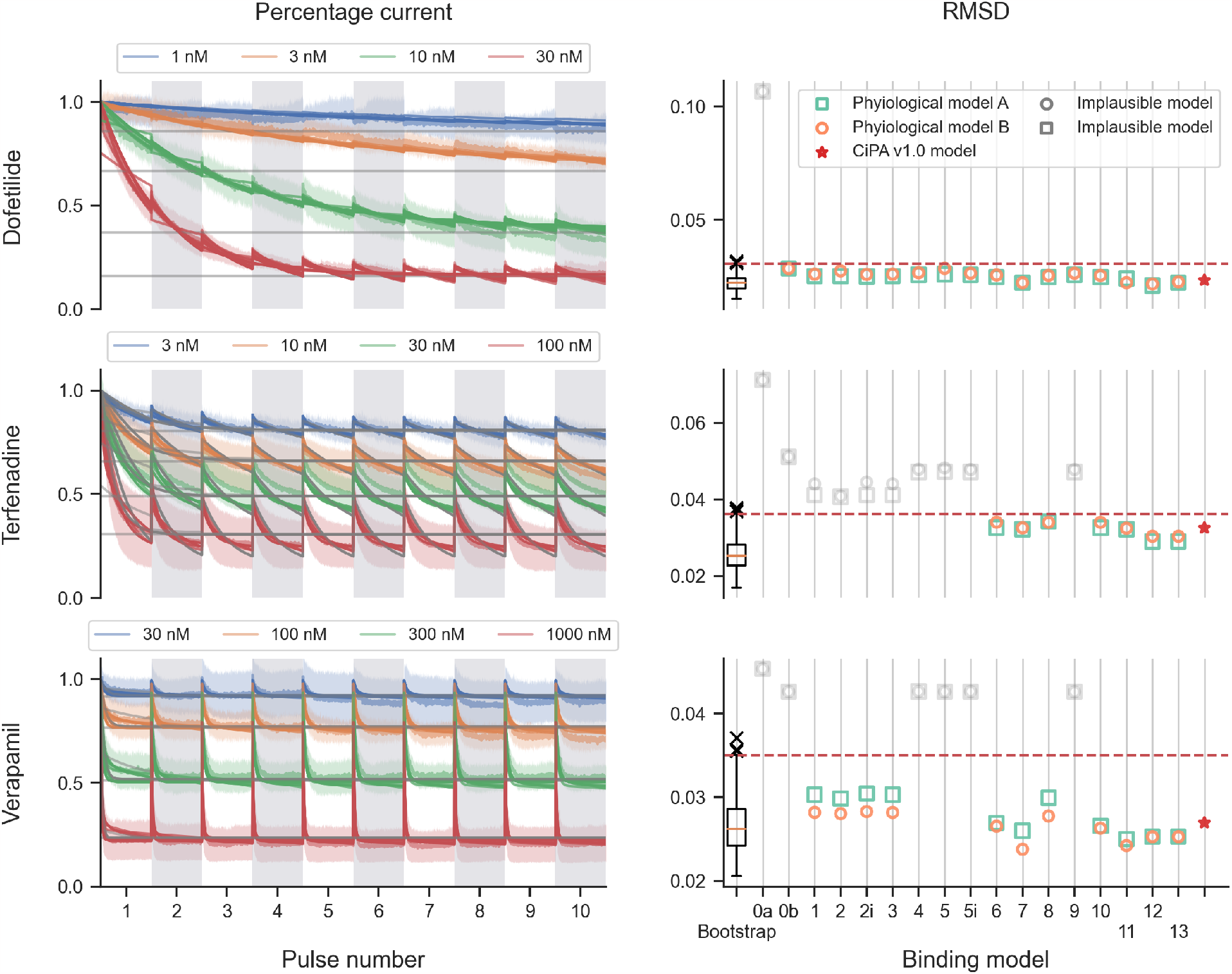
Calibration of the binding models and their RMSDs for three example compounds: dofetilide (**top row**), terfenadine (**middle row**), and verapamil (**bottom row**). **(Left column)** Shows the percentage current of the data (transparent lines) and the calibrated binding models (solid lines) for all four concentrations used during calibration. **(Right column)** Shows the RMSD of all models compared to the RMSD of the bootstrap samples of the data (box-plot) and the reference model, CiPA v1.0 (red star, Li et al., 2017). Both physiological models A (green squares) and B (orange circles) are shown for comparison. The horizontal red dashed line indicates the maximum RMSD of bootstrapped samples of the data Grey lines/markers on each panel are the binding models that are ruled out through the RMSD comparison (above the red dashed line)—implausible models.

For dofetilide, a trapped drug (Milnes et al., 2010), *all* binding models except the steady-state conductance scaling model (Model 0a) were able to fit the calibration data, and so we cannot unpick the binding mechanisms of this drug with these data. The four grey horizontal lines in Figure 3 (left) are predictions of Model 0a, the only implausible model for dofetilide. Model 0a was the only model that showed no dynamics in the percentage current plots as, by definition, it scales only the conductance of the model so its effect on *I*_Kr_ must be constant over time. Another model that lacked in the drug dynamics was Model 0b, the all-state binding model, which marginally passed the RMSD check. Model 0b modelled the drug effect with the *b* gate in Eq. (2) that is independent of channel state/voltage, producing a single exponential decay over time—even during the long resting potential (−80 mV) of CiPA-Milnes’ protocol (although not shown) when the channels were in the closed state(s), resulting in the drop in current between pulses, when the data show (if anything) a slight restoration of current between pulses.

The other two example compounds in Figure 3, terfenadine and verapamil, had a weaker to no trapping tendency compared to dofetilide; that is, current recovers significantly between 0 mV pulses, suggesting unbinding rather than trapping at −80 mV. Interestingly, more binding models failed to fit to the experimental data of these two drugs. For terfenadine, a weakly-trapped drug (Yang et al., 1995; Kamiya et al., 2008; Stork et al., 2007) with slow binding rates (Li et al., 2017), only Models 7, 8 (variants of the independent open and inactivated binding model) and Models 6, 10–13 (variants of *flexible*-trapping models) were able to explain its drug effect as well as the bootstrap samples of the data, although a few more non-trapping models were merely marginally ruled out. For verapamil, a non-trapped drug (Zhang et al., 1999), our approach successfully disqualified *all simple*-trapping models (Models 4, 5, 5i, and 9), as well as the conductance blocking and the all-state binding models (Models 0a and 0b). However, it left the non-trapping models (Models 1–3, 7, and 8) and the flexible-trapping models (Models 6, 10–13) as plausible candidates for the binding mechanism.

Figure 4 presents a summary of the binding models with physiological model A which gave plausible fits to the experimental data (shown in green); a summary for those with physiological model B are provided in Supplementary Figures, the results were very similar with only minor differences for cisapride, ibutilide and loratadine. We observed no obvious pattern between the proarrhythmic risk of drugs and the type of binding models ruled implausible. A few more non-trapped drugs were identified simply based on the ruling out of the simple-trapping models (Models 4, 5, 5i, and 9) as in the case of verapamil (Figure 3), such as cisapride (Milnes et al., 2010) and droperidol (Stork et al., 2007; Windisch et al., 2011). For most of the drugs, multiple binding models were able to explain the observed drug effects using the experimental data collected through CiPA-Milnes’ protocol.

**Figure 4:**
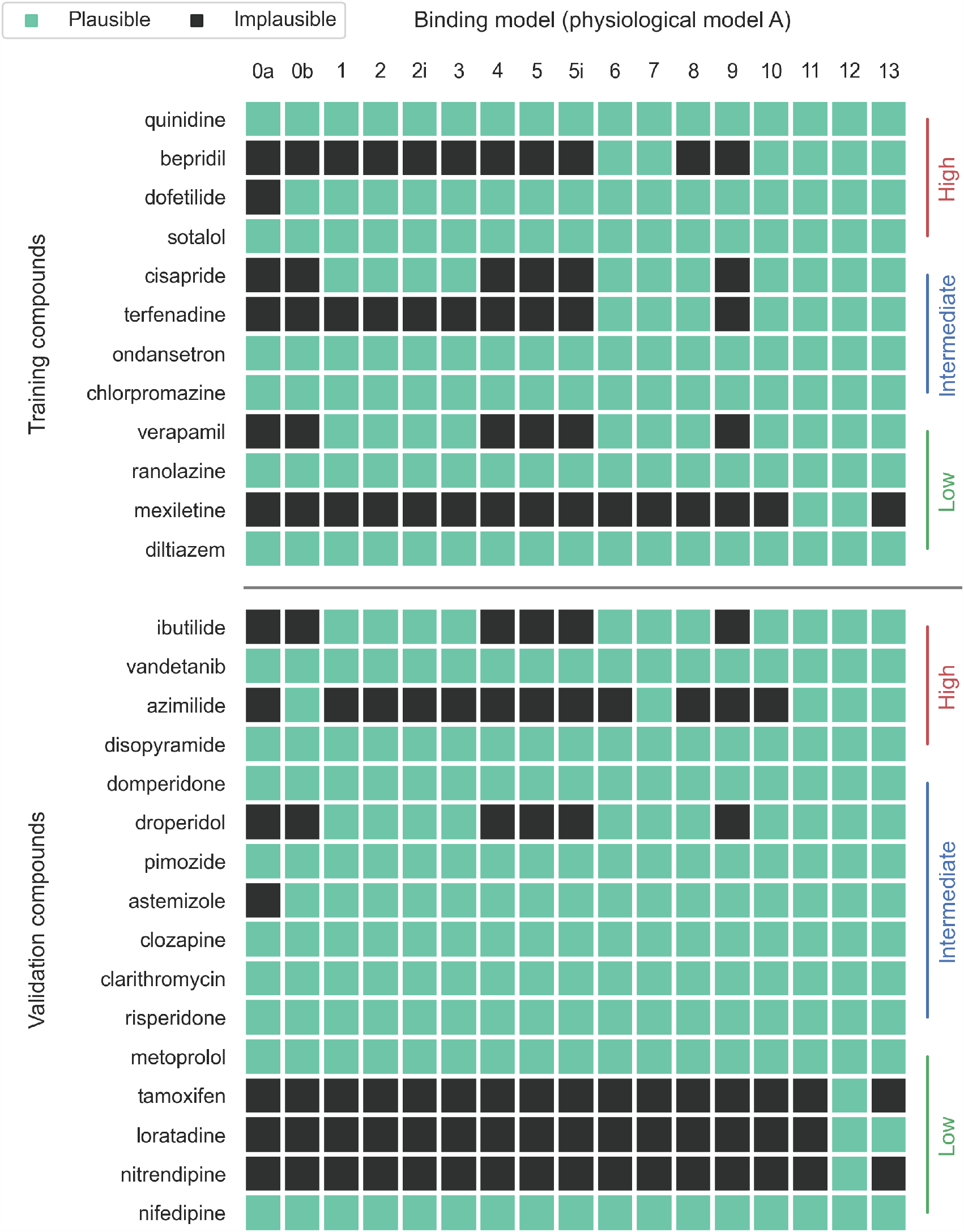
Summary of the selected binding models with physiological model A for all compounds through the RMSD comparison. A binding model (column) is considered to be appropriate for a compound (row)—a plausible model—if coloured in green, where the RMSD of the model to the averaged data is either smaller than the RMSD of the bootstrap samples of data or similar to the CiPA v1.0 model to the averaged data. Model 12 is identical to the pharamcological binding component in the CiPA v1.0 model. Drugs are sorted according to the training and validation lists, and their proarrhythmic risks.

### 3.2 Binding rates can be strongly dependent on binding mechanism

Studying drug binding kinetics, such as the binding and unbinding rates, is important for understanding the drug effects on the channels and predicting behaviour in new situations. Figure 5 shows inferred binding rate parameters *k*_on_, unbinding rates *k*_off_, and the Hill coefficients *n* of the calibrated binding models. All binding models have *k*_on_, *k*_off_, and *n*, apart from the conductance scaling model (Model 0a) which has only two parameters IC_50_ and *n* (Table 1). Models 7–10 have independent binding rates for the open and inactivated states, giving two *k*_on_ shown as empty (for open) and filled (for inactivated) markers; only Model 7 has two (free) *k*_off_, shown in the same way, as Models 8–10 have closed-loop states which reduce one degree of freedom due to microscopic reversibility (Figure 2). The CiPA v1.0 model is shown as red stars; the two physiological models, A and B, are shown together as squares and circles, respectively, and the observed results remained similar regardless of the physiological model. The results for all the remaining compounds are shown in Supplementary Figures.

**Figure 5:**
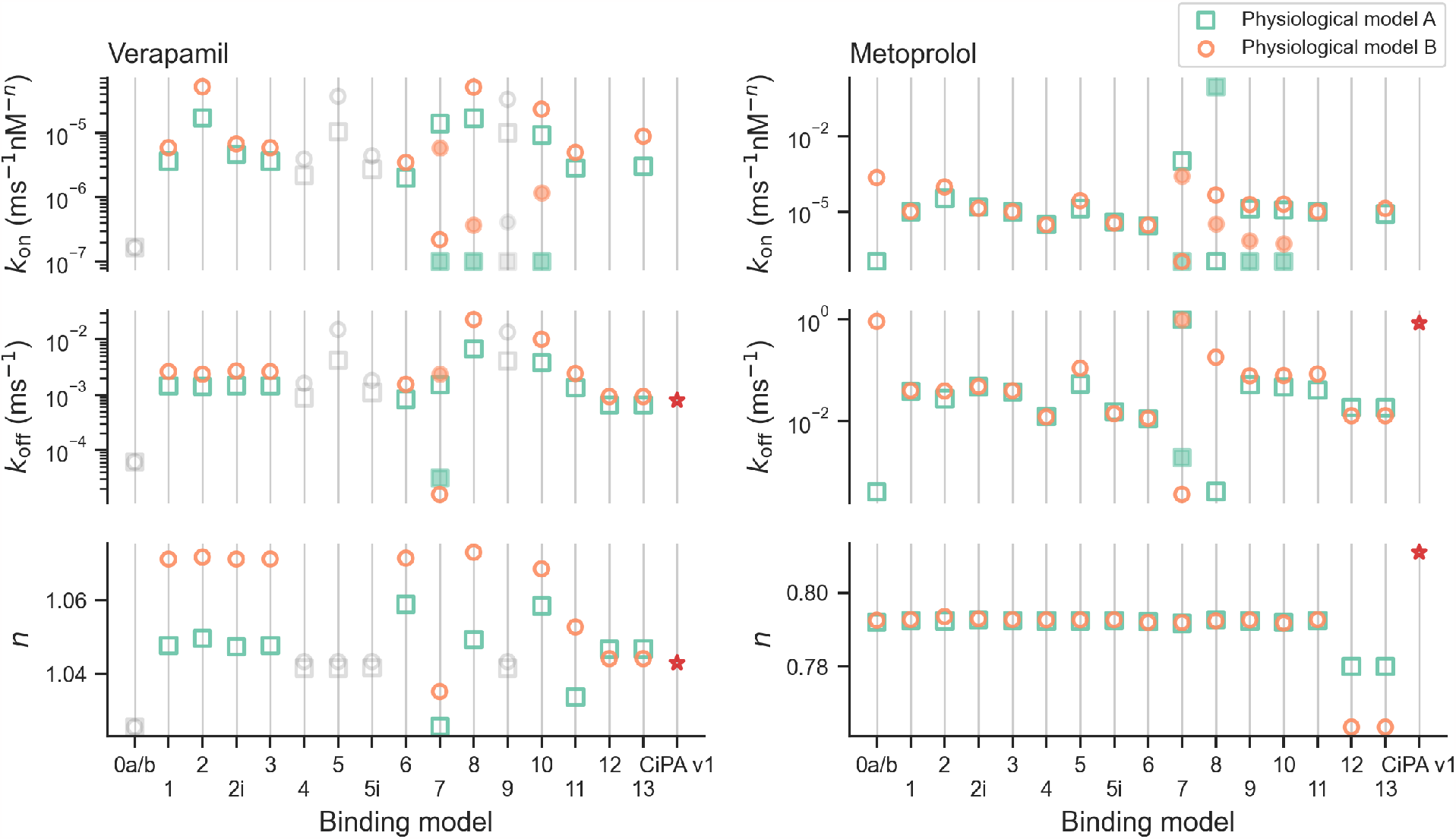
Binding rate parameters *k*_on_ (**top row**), unbinding rates *k*_off_ (**middle row**), and the Hill coefficients *n* (**bottom row**) of the calibrated binding models for two example compounds: verapamil (**left column**) and metoprolol (**right column**). Both physiological models A (green squares) and B (orange circles) are shown for comparison. Models 7–9 have independent binding and unbinding rates for open and inactivated states; filled squares/circles are the rates for the inactivated states. The models in grey are implausible models ruled out through the RMSD comparison. Model 12 is identical to the pharmacological binding component in the CiPA v1.0 model (red star). 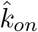 is not shown for Model 12 and CiPA v1 due to different units.

Taking verapamil as an example, the inferred binding rate parameter *k*_on_ was similar for most of the successfully calibrated binding models (Figure 3). The two binding rate parameters *k*_on,O_ (empty marker) and *k*_on,I_ (filled marker) for the plausible models—Models 7, 8, and 10—were similar too, and their average roughly equalled *k*_on_ of the other models. The similarity suggested that the independence assumption may be superfluous in this case, because if the two inferred rates were the same, the models would be equivalent to those without the extra degree(s) of freedom (Models 1, 3, and 6). Similarly, the unbinding rates *k*_off_ were similar for all binding models, except the *k*_off,O_ (empty marker) and *k*_off,I_ (filled marker) in Model 6; all of the inferred Hill coefficients *n* were within the range of 1.3–1.8. It is worth noting that the *k*_off_ of Model 12 and the CiPA v1.0 model (the same binding model structure but under different calibration schemes) shared similar inferred values for verapamil but their 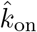 were not shown due to different units.

For metoprolol, the inferred *k*_on_ were heavily binding-model-dependent, with a coefficient of variation (defined as the ratio of the standard deviation to the mean, for the plausible models) of 607 %, as compared to 141 % for verapamil. The other two parameters *k*_off_ and *n* were similar across the calibrated binding models. Unlike verapamil, the inferred *k*_off_ of Model 12 and the CiPA v1.0 model were inconsistent, using the same set of experimental data but with different calibration schemes; see also mexiletine in Supplementary Figures. The differences in the inferred parameters may raise questions about the calibration data and the complexity of the model structure used, leading to potential parameter unidentifiability issues (Whittaker et al., 2020, see also Discussion).

### 3.3 *q*_net_ with different binding mechanisms diverges at higher drug concentration

Thus far, we compared the binding models of hERG when calibrated to voltage-clamp experimental data for various compounds. Here, we show the results of our investigation on the impacts of these binding models on APs and risk simulations (§ 2.4), whilst accounting for multi-channel effects, following Li et al. (2019). Figure 6 shows the metric *q*_net_ at various *C*_max_ levels for all binding models, with implausible models shown with dotted lines. The torsade metric decision boundaries for the low- (green), intermediate- (blue) and high- (red) risk categories in Li et al. (2019) are indicated as dashed horizontal lines for reference. Various degrees of *q*_net_ spread were observed across the binding models for different compounds.

**Figure 6:**
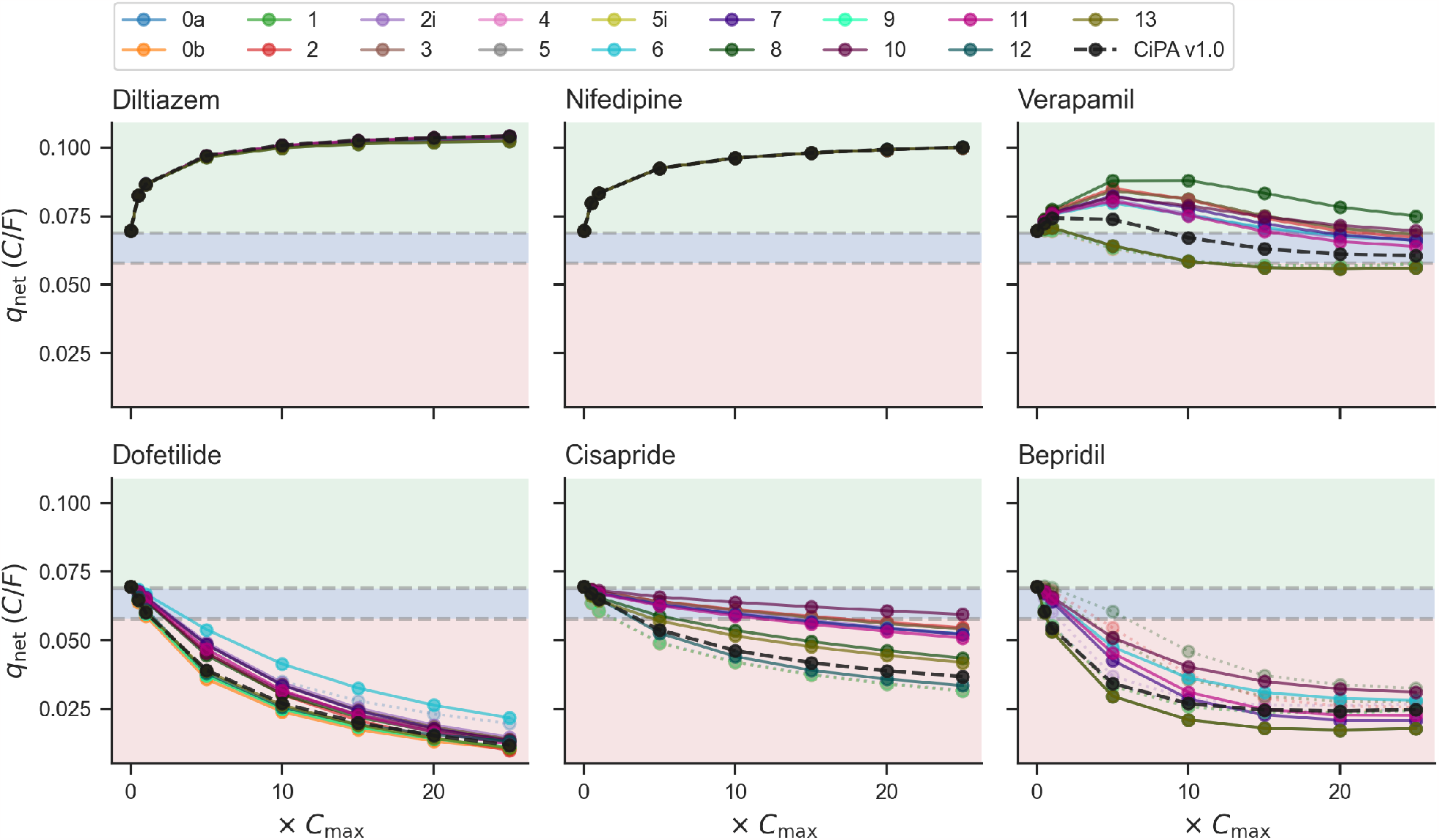
*q*_net_, the net charge carried by currents active in plateau and repolarisation over one beat of AP (*I*_Kr_, *I*_NaL_, *I*_CaL_, *I*_to_, *I*_Ks_, and *I*_K1_) at various multiples of *C*_max_ for all binding models with six example drugs. Here, diltiazem, nifedipine, verapamil, dofetilide, cisapride, and bepridil are shown, revealing a spectrum of behaviours from the binding models. Dashed horizontal lines indicate the decision boundaries for the low- (green), intermediate- (blue) and high- (red) risk categories for the torsade metric score (average *q*_net_ at 1–4× *C*_max_) in Li et al. (2019). The implausible models are shown as transparent dotted lines, and the CiPA v1.0 model is shown as black dashed lines.

For example, diltiazem is a strong *I*_CaL_ blocker relative to its *I*_Kr_ effects, therefore the drug effect on APs measured through *q*_net_ reflects mostly the drug block of *I*_CaL_ and the effects of different *I*_Kr_ binding models are insignificant. Nifedipine and verapamil are also multi-channel blockers of *I*_CaL_ and *I*_Kr_, with similar block levels for each current. However, the uncertainty in their resulting *q*_net_ predictions were drastically different—nifedipine showed tight *q*_net_ predictions across the binding models, whilst verapamil produced a wide spread. Figure 6 (bottom row) also shows three more examples: dofetilide, cisapride, and bepridil. These compounds are almost pure *I*_Kr_ channel blockers at these concentrations. Again, they showed inconsistent spread of *q*_net_ predictions from the *plausible* binding models, demonstrating the importance of identifying the correct binding mechanisms or at least narrowing down the possibilities.

Furthermore, in general, we also observed an increasing spread of the *q*_net_ predictions from plausible binding models at higher *C*_max_ levels; verapamil is one of the most obvious examples. This phenomenon showed the differences between the binding mechanisms were amplified with higher concentrations of the drug, and at the top concentrations verapamil spanned all risk categories. However, the torsade metric uses only 1–4× *C*_max_ of *q*_net_—the metric that was used to train the classification model (ordinal logistic regression model) for producing the decision boundaries of the risk categories, and we examine risk predictions in this range in the next section.

### 3.4 Binding mechanisms can result in substantial uncertainty in torsade risk

Figure 7 (left) shows the torsade metric predictions of all binding models with physiological model A for all compounds. The drugs are sorted according to their proarrhythmic risk categories and are split into training and validation lists (Li et al., 2019); the same decision boundaries as Figure 6 are shown as dashed vertical lines, for the low- (green), intermediate- (blue) and high- (red) risk categories.

**Figure 7:**
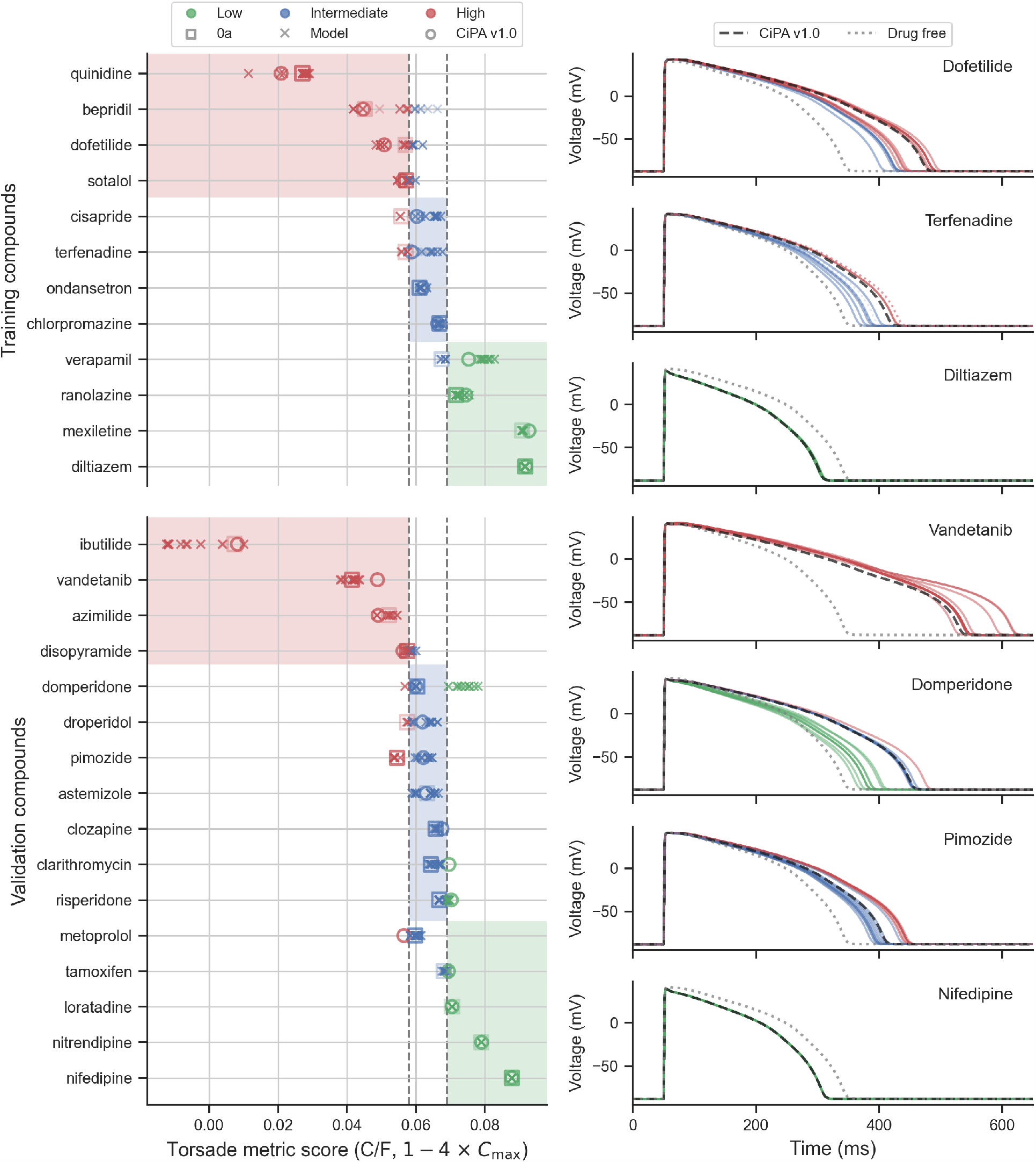
The torsade metric predictions, the mean *q*_net_ of 1–4× *C*_max_, of all binding models with physiological model A for all compounds, and the corresponding AP predictions for seven example drugs. **(Left)** Drugs are sorted according to their proarrhythmic risk categories. Dashed vertical lines indicate the CiPA v1.0 decision boundaries for the low- (green), intermediate- (blue) and high- (red) risk categories (Li et al., 2019). The CiPA v1.0 model is shown as circles, and the conductance scaling model, Model 0a, is shown as squares. **(Right)** The AP predictions of dofetilide, terfenadine, diltiazem, vandetanib, domperidone, pimozide, and nifedipine at 4× *C*_max_ are shown, revealing a range of different behaviours from the binding models, indicated with the same colour code as shown on the left. The implausible models are shown as transparent dotted lines, and the CiPA v1.0 model is shown as black dashed lines. The drug-free (control conditions) model is shown as grey dotted lines.

The torsade metric predictions for the same compound resulted in a large variation—indicating high uncertainty in the proarrhythmic risk prediction—due to different binding mechanisms. However, it is worth emphasising that these plausible binding models were able to explain the observed experimental data of the compounds either better than the bootstrap samples of the data or as well as the CiPA v1.0 model, therefore the acquired experimental data were not able to resolve the resulting uncertainty. The reference model, CiPA v1.0, is shown as circles, and the conductance scaling model, Model 0a, is shown as squares; unexpectedly, Model 0a, which had the worst performance during the calibration process (Figure 4) and was ruled out in most cases, did not cause any obvious outlying torsade metric prediction (cf. the extreme RMSD values of Model 0a in Figure 3), and in fact it was usually not the one giving the predictions at the extremes, showing the nonlinearity in the relationship between the calibration and the torsade metric prediction. Also, interestingly, the degree of variation tended to be larger for drugs in higher proarrhythmic risk categories, for both training and validation drugs, perhaps because hERG block was more dramatic in these compounds (see Discussion).

The corresponding AP predictions for some of the drugs at 4× *C*_max_ are shown in Figure 7 (right). In general, as expected, a strong AP prolongation—longer AP duration—correlated with a high torsade metric risk category: the shortest AP duration for the low-risk category in green, followed by intermediate in blue, and then the longest AP for high risk in red. The plausible binding models can result in metric predictions with high uncertainty, even spanning multiple risk categories; for example, domperidone predictions span all three risk categories.

### 3.5 The effect of hERG physiological model on risk classification is subtle

Finally, we compared the effects of the choice of the *I*_Kr_ physiological model (Figure 1) in predicting the torsade risk classes. The “O’Hara-Rudy CiPA v1.0” model (Dutta et al., 2017) had hERG physiological model A replaced with physiological model B (solid lines, Lei et al., 2019b,a), the *I*_Kr_ maximum conductance was re-calibrated to match the APD_90_ of the CiPA v1.0 AP model (dashed lines, Li et al., 2017), as shown in Figure 8A (blue lines). The *I*_Kr_ within the two AP models, shown as orange lines, reveals differences in the dynamics of the two hERG physiological models. Although the total charge carried by the two *I*_Kr_ models is similar (0.190 C/F and 0.144 C/F for A and B, respectively), physiological model B has a smaller current during the early phase of the AP—depolarisation to plateau—and plays a more important role during repolarisation. The two *I*_Kr_ physiological models show a similar overall transition between the states (Figure 8A): starting from mainly the closed state(s) to the inactivated state(s) during depolarisation/plateau before occupying the open state and back to the closed state(s).

**Figure 8:**
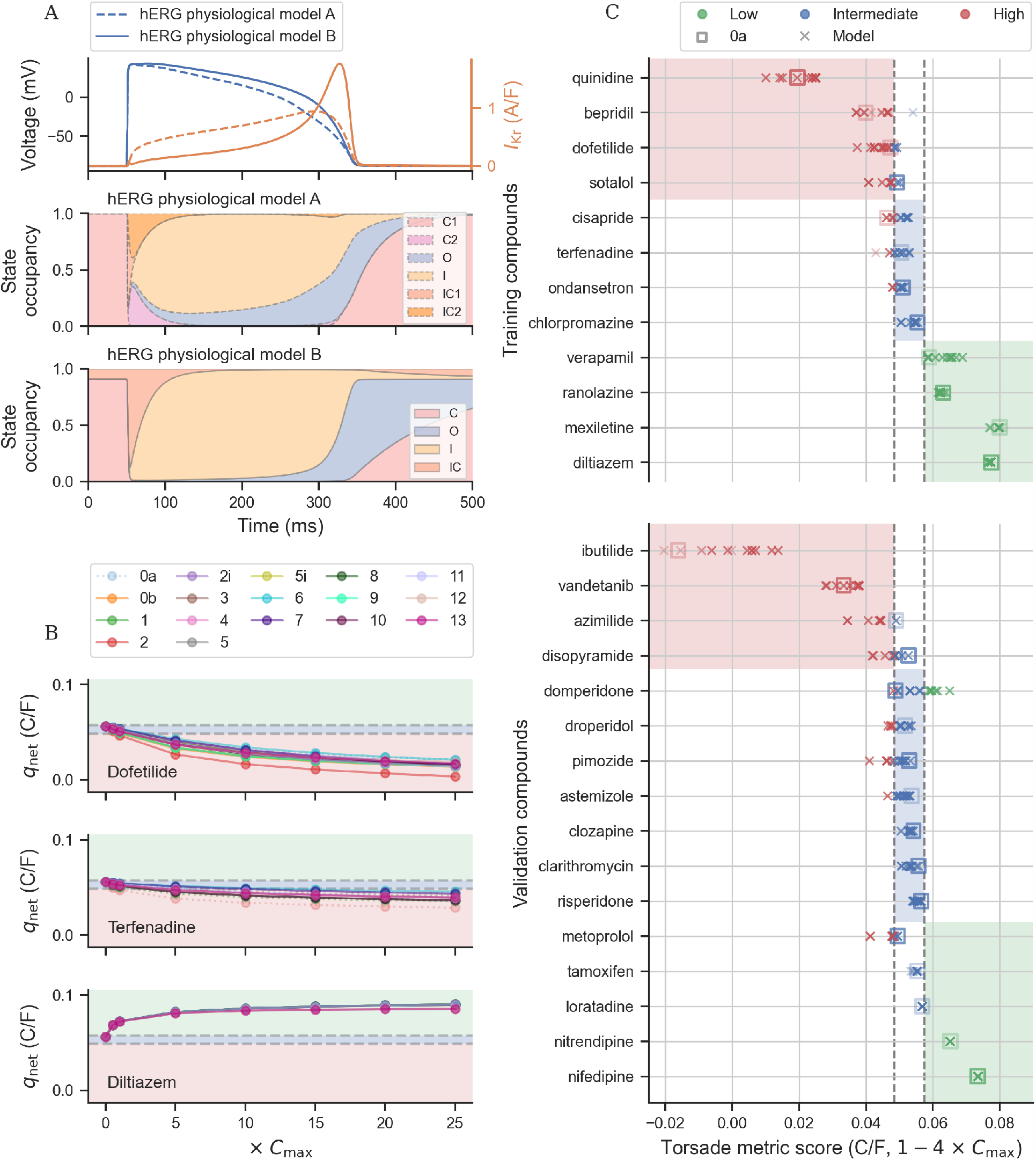
Effects of hERG physiological model B in predicting the torsade metric. **(A)** Comparison of AP, *I*_Kr_, and state occupancy using the two hERG physiological models A (dashed lines) and B (solid lines). **(B)** The *q*_net_ metric at different *C*_max_ levels for all binding models with physiological model B for dofetilide, terfenadine, and diltiazem. **(C)** The torsade metric predictions of all binding models with physiological model B for all compounds. Dashed vertical/horizontal lines indicate the new decision boundaries for the low- (green), intermediate- (blue) and high- (red) risk categories, using an ordinal logistic regression model.

Training the ordinal logistic classification model to the torsade metric of the training drugs for all the *plausible* binding models adjusted the risk category thresholds to be 0.0575 C/F and 0.0484 C/F for separating the low-risk category from intermediate/high and the high-risk category from intermediate/low, respectively. The shift in the boundaries was consistent with the change in the net charge carried by the physiological model B (and alteration to AP shape which also affects other currents) which resulted in a new control value of *q*_net_. The two new decision boundaries are indicated as dashed lines in Figures 8B and C. Replacing the physiological model of *I*_Kr_ still produced a similar trend for the drug risk categories as we saw in Figure 7. Although some of the drugs, such as bepridil, were clustered closer to the new decision boundaries. This could result from the new *I*_Kr_ model, the choice of calibration protocol, as well as the choice of the metric, all of which were designed for CiPA v1.0.

## 4 Discussion

In this study, we have designed a set of pharmacological binding models for the hERG channel. After selecting a subset of plausible binding models through calibration to the voltage-clamp electrophysiology data under CiPA-Milnes’ protocol, we have compared their effects, as well as the effects of having different hERG physiological models, on proarrhythmic risk predictions.

Our pharmacological binding models of hERG accounted for most of the plausible mechanisms by which a compound might bind to the hERG channel. The choice of these models has covered some of the literature binding models of hERG. For example, our simple binding models (Models 2, 2i, and 7) and trapping models (Models 5, 5i, and 9) are similar to models referred to as “unstuck” and “stuck”, respectively, in Gomis-Tena et al. (2020). Models 0a, 7 and 11 were used in Windley et al. (2016) to study the effects of compounds such as cisapride.

In Figures 3 and 4, we demonstrated that our approach can currently be used to distinguish some of the simpler binding mechanisms for some compounds, such as terfenadine and verapamil, consistent with the literature. It was also able to highlight certain similarities between some compounds such as bepridil (Kamiya et al., 2006; Pareja et al., 2013) and terfenadine (Stork et al., 2007) where only flexible-trapping models were deemed plausible to explain the observed data; indeed, studies with specifically designed voltage protocols consider these two compounds to be trapped slow-binders (Kamiya et al., 2008). Also, our approach was able to highlight compounds, such as tamoxifen, loratadine, and nitrendipine, where none of the models (not even the CiPA 1.0 model) were able to fit the data satisfactorily. On closer inspection (Supplementary Figures) the data of loratadine, and nitrendipine showed a slight *increase* of the (percentage) current over time during the 0 mV pulses, raising potential data quality issues (Lei et al., 2020a) or the need for methods to account for inadequacy of the models (Lei et al., 2020b) and/or new (un)binding mechanisms to explain this observation; whilst for tamoxifen, there was a more obvious data quality issue for one of the concentrations. In Supplementary Figures, we also included the results of fitting all the binding models whilst assuming the Hill coefficient (number of binding sites) to be *n* = 1. However, in this case, most of the binding models failed to fit to many of the compounds—being classified as implausible models—suggesting the importance of the extra degree of freedom provided by the Hill coefficient in explaining these observations, although we do not eliminate the possibility of this having been caused by some experimental artefact resulting in a drift/rundown that was mistaken for non-saturating or more quickly/slowly saturating hERG block. Ideally, we would like the calibration data to contain enough information to rule out as many binding models as possible. We can use the approach presented in this study to objectively and quantitatively describe a compound’s binding mechanism instead of qualitatively classifying the compound as “trapped”, “binding to open state”, etc., as well as to predict future behaviour in new situations, hERG mutations in patients, etc. However, we observed that the data elicited with CiPA-Milnes protocol, which was originally designed to differentiate between trapped and non-trapped compounds (Milnes et al., 2010), were *not* able to distinguish between the possible binding mechanisms, or indeed whether trapping occurs, for all compounds (Figure 3). It therefore suggests the need for designing better, richer experimental protocols for model calibration and selection, for instance gathering data on block onset at different voltages (Lee et al., 2019; Gomis-Tena et al., 2020), overcoming the difficulties in measuring some timecourse of block (Windley et al., 2017), whilst the protocols may also need to be suitable for fast- and slow-binding compounds that bind via multiple mechanisms.

The model structure is not only useful for identifying the binding mechanisms; the inferred model parameters, such as binding/unbinding rates, for the *plausible* pharmacological binding model(s) can be used as a proxy to quantitatively assess the binding behaviour and dynamics. However, we showed that, with the limitations of the calibration data, not all plausible models recover the same binding and unbinding rates, although their ratios, the dissociation constants *k*_off_ */k*_on_ of the models, were more consistent (Figure 5 and Supplementary Figures). This leads to a more complicated interpretation of the binding properties. Without being able to identify the correct binding mechanism(s), we would not be able to study the true binding/unbinding rates of the compound. Moreover, for more complex pharmacological models, the inferred parameters may even be subject to the calibration scheme and procedure: e.g. the inferred unbinding rate *k*_off_ of Model 12 and CiPA v1.0 were inconsistent (Figure 5), although our Model 12 gives lower RMSDs (Supplementary Figures). The different optimal parameters in CiPA v1.0 versus Model 12 fits could be due to the use of a different objective function—a tailored weighted sum of squares of residuals, and based on fewer optimisation runs which could make local rather than global optima more likely (Li et al., 2017). We note that this is not the same unidentifiability issue between, in our notation, 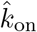 and EC_50_ discussed in Li et al. (2019), suggesting this is a difficult optimisation problem. Our findings emphasise the need for designing better experiments when parameterising complex models (Whittaker et al., 2020).

In Figure 4, we observed that Model 0a was not classified as a plausible model for most of the compounds, and in fact the goodness of fits were inadequate and poor (Figure 3). Yet, the torsade metric predictions by Model 0a were not dissimilar to the other models (Figure 7, see also Mistry, 2019; Han et al., 2019), which was likely due to the difference between using transient data (Milnes’ protocol) and predicting steady-state (*q*_net_/torsade metric) conditions (Farm et al., 2023). However, we believe the torsade metric predictions of Model 0a were acceptable only because of the steady-state nature of the metric. If Model 0a was used to predict transient APs under certain changes of conditions, then the trapping effects would be neglected.

In Figure 7, we also noticed that the degree of variation tended to be larger for drugs in higher proarrhythmic risk categories. This phenomenon was thought to be due to the multi-channel effects, which either compensate for the effects of different hERG binding mechanisms or make them insignificant. However, if such an analysis was applied to other types of current, such as *I*_CaL_, etc., it may result in a similar level of uncertainty in their binding mechanism, leading to a higher level of uncertainty in the risk predictions than we observed (Figure 7) for compounds that mainly block e.g. *I*_CaL_ (diltiazem and nifedipine). Nonetheless, such an observation also implies that it is likely to be more important to determine the correct hERG binding mechanism when predicting drugs with higher proarrhythmic risk. Also, given some of the risk prediction spanned multiple risk categories, we advocate efforts to reduce the uncertainty. Overall, the results highlight the importance of the details of hERG binding mechanisms.

Furthermore, we have studied the effects of hERG physiological models on proarrhythmic risk predictions (Figures 7 and 8). To choose between the two hERG physiological models (Figure 1), since the calibration data were gathered using cell lines expressing hERG1a, we expect that physiological model B, designed to be a hERG1a model, could capture the drug effects better. Yet, since the CiPA v1.0 model and the torsade metric were optimised and designed with physiological model A, which was supposed to capture the native *I*_Kr_ with hERG1a/1b channels (Jones et al., 2004), we may expect it to predict these risk metrics better. We could fit the hERG1a data with physiological model B to obtain the binding parameters, then apply these binding parameters to physiological model A to predict changes to native *I*_Kr_ and clinical risk. However, it would perhaps be more convincing to directly perform voltage-clamp experiments on hERG1a/1b cell lines (Ríos-Pérez et al., 2021) when calibrating these pharmacological binding models, as compounds with differing affinities for hERG1a and 1b have been observed (Abi-Gerges et al., 2011).

Finally, to our knowledge, this is the first study that attempts to address the question of whether we need a complex pharmacological model that attempts to nest most/all of these binding mechanisms or a set of multiple possible simpler pharmacological models to perform better proarrhythmic risk predictions. In theory, if parameter unidentifiability were not an issue, the two wings should arrive at the same conclusion—for example, when modelling an untrapped compound, the transition rates to the trapping component of the complex model would approach zero, and the non-trapping (simpler) model would be selected as the only plausible model. However, it may be better to use simpler models given the difficulty of eliciting information-rich data for calibrating complex binding models leading to potential parameter unidentifiability issues (Whittaker et al., 2020), and the inevitable presence of some model discrepancy (Lei et al., 2020b) and residual experimental artefacts (Lei et al., 2020a). Pragmatically, based on our results, we would suggest to select and use all the plausible simpler models for prediction as demonstrated here—a type of ensemble model prediction which provides an estimate of uncertainty due to model discrepancy (Murphy et al., 2007; Tebaldi and Knutti, 2007; Parker, 2013). In conclusion, this study has developed an approach to analyse a set of possible pharmacological small molecule binding models of hERG that is effective in assessing their impacts, as well as the impact of different physiological *I*_Kr_ models, on the proarrhythmic risk predictions. Determining the details of binding mechanisms, perhaps through the design of an improved calibration protocol, is crucial for mitigating the induced, substantial uncertainty in risk predictions for some compounds.

## Data Availability Statement

The source code and data that support the findings of this study are openly available in GitHub at https://github.com/chonlei/hERG-binding-mechanisms [to be archived on Zenodo and assigned a DOI upon acceptance].

## Author Contribution Statement

CLL, DGW and GRM conceptualised and designed the study. CLL and DGW wrote the code. CLL performed the simulations and data analysis, and designed the visualisation and generated the results figures. CLL and GRM wrote the manuscript. All authors revised and approved the final version of the manuscript.

## Acknowledgements

We would like to thank a number of people who have helped to form our ideas on potential modes of hERG binding: Adam Hill, Jamie Vandenberg, Randall Rasmusson, Jules Hancox, Tom Claydon, Gail Robertson, Erick Rios Perez, Ken Wang, Derek Leishman & Zhihua Li.

