## Supplementary Figures for "The impact of uncertainty in hERG binding mechanism on *in silico* predictions of drug-induced proarrhythmic risk"

Figures on following pages.

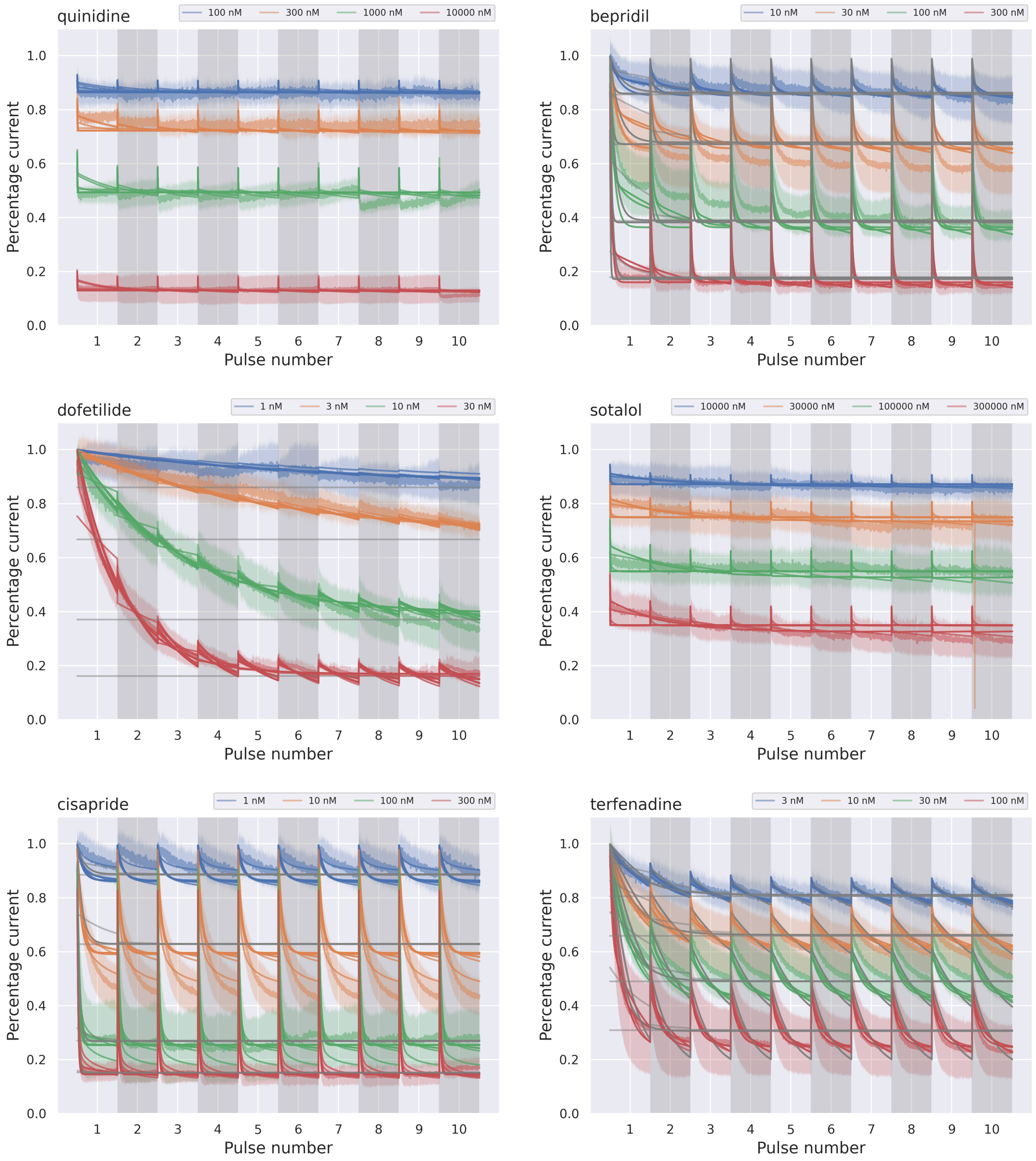


Figure S1: The percentage current of the data (transparent lines) and the calibrated binding models (solid lines) for all four concentrations used during calibration. Grey lines are the binding models that are ruled out through the RMSD comparison.

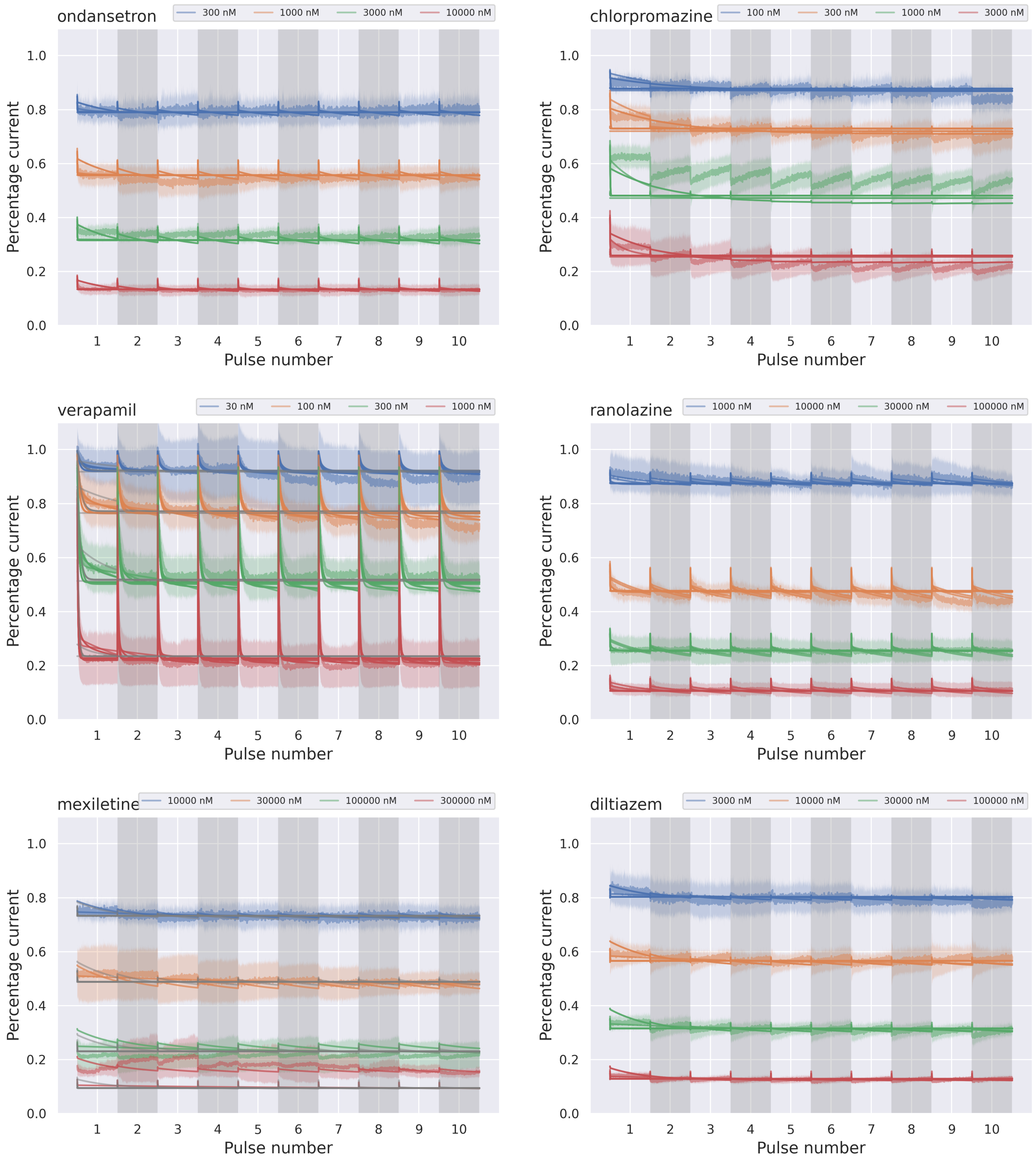


Figure S2: The percentage current of the data (transparent lines) and the calibrated binding models (solid lines) for all four concentrations used during calibration. Grey lines are the binding models that are ruled out through the RMSD comparison.

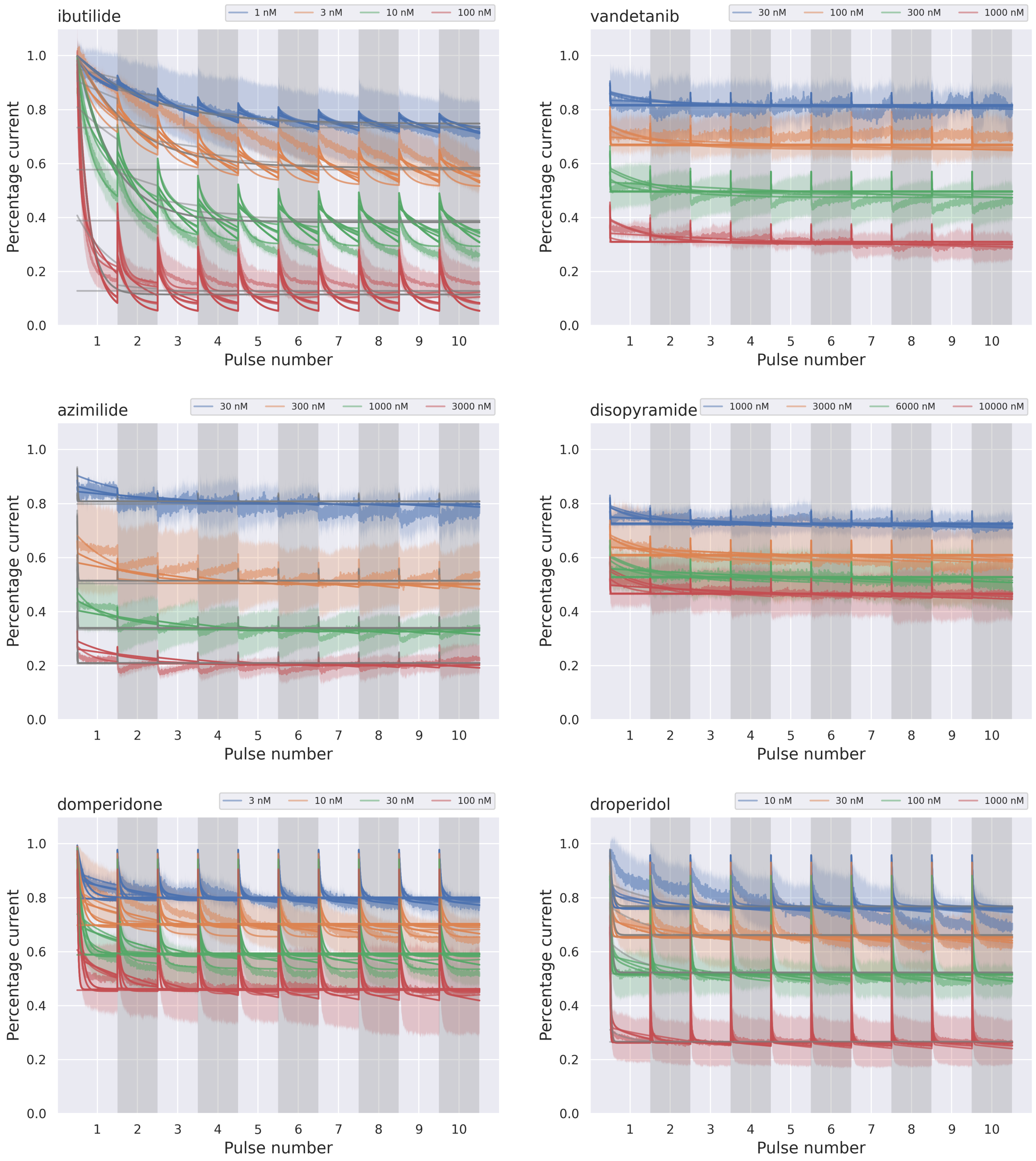


Figure S3: The percentage current of the data (transparent lines) and the calibrated binding models (solid lines) for all four concentrations used during calibration. Grey lines are the binding models that are ruled out through the RMSD comparison.

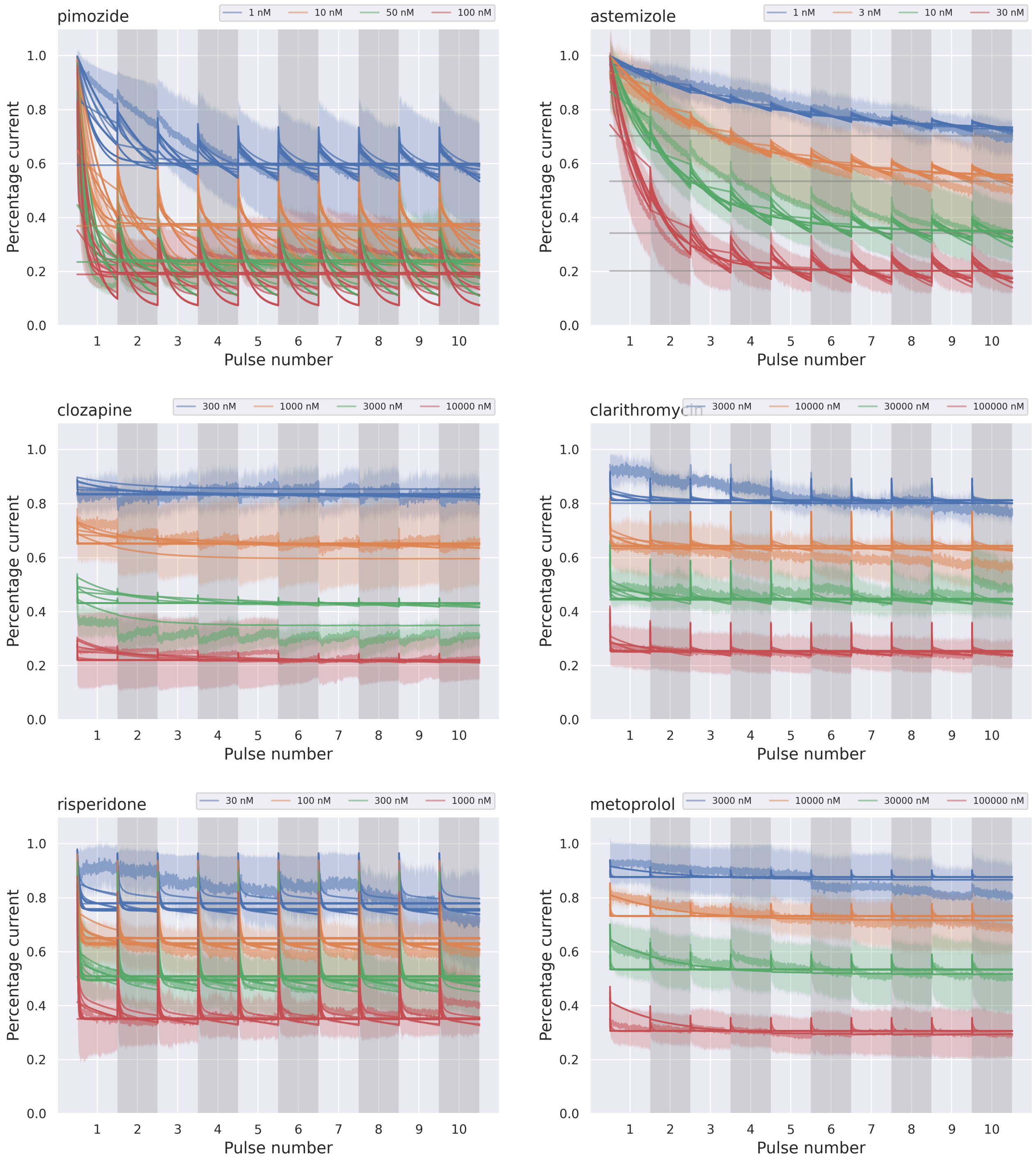


Figure S4: The percentage current of the data (transparent lines) and the calibrated binding models (solid lines) for all four concentrations used during calibration. Grey lines are the binding models that are ruled out through the RMSD comparison.

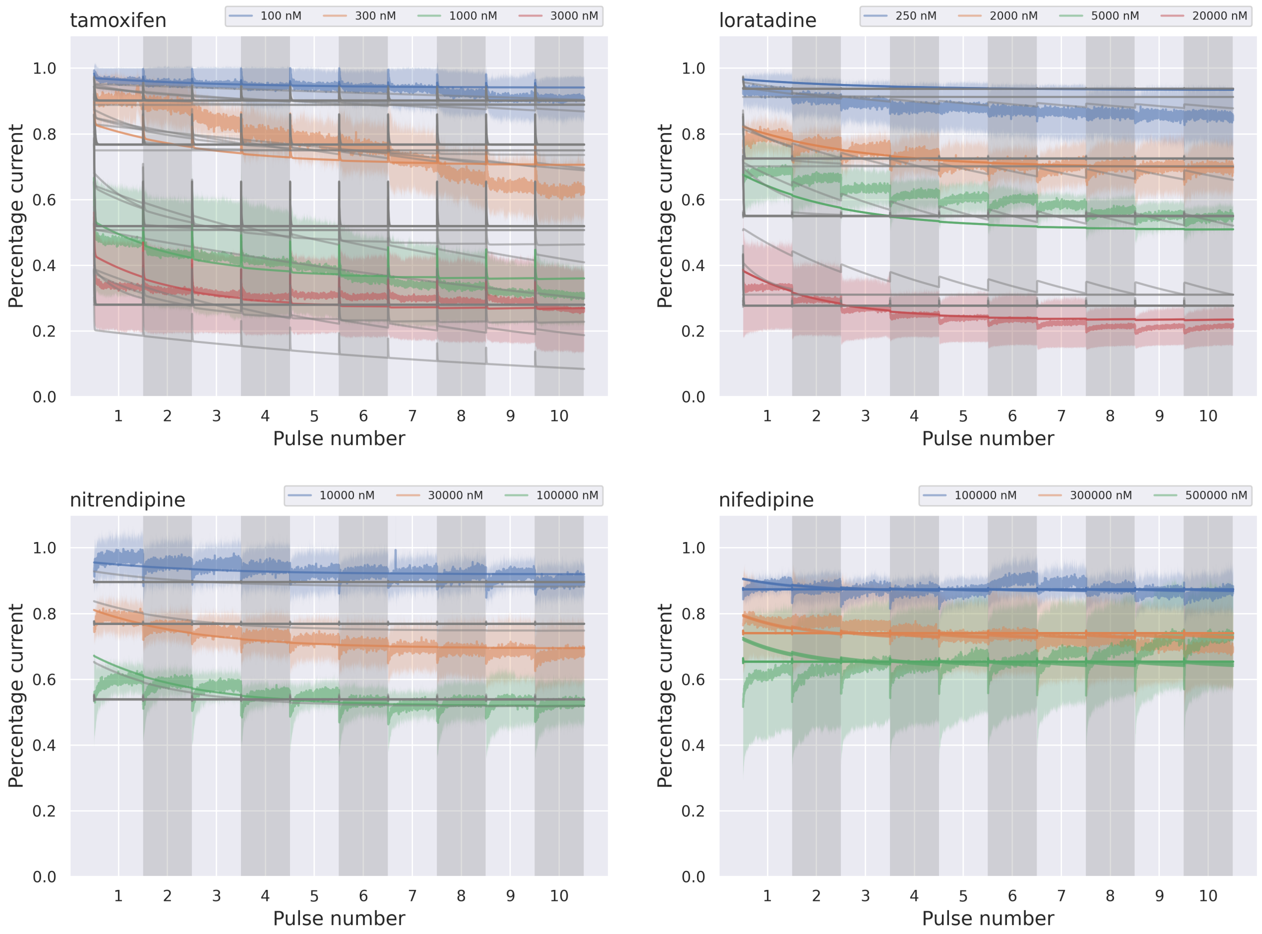


Figure S5: The percentage current of the data (transparent lines) and the calibrated binding models (solid lines) for all four concentrations used during calibration. Grey lines are the binding models that are ruled out through the RMSD comparison.

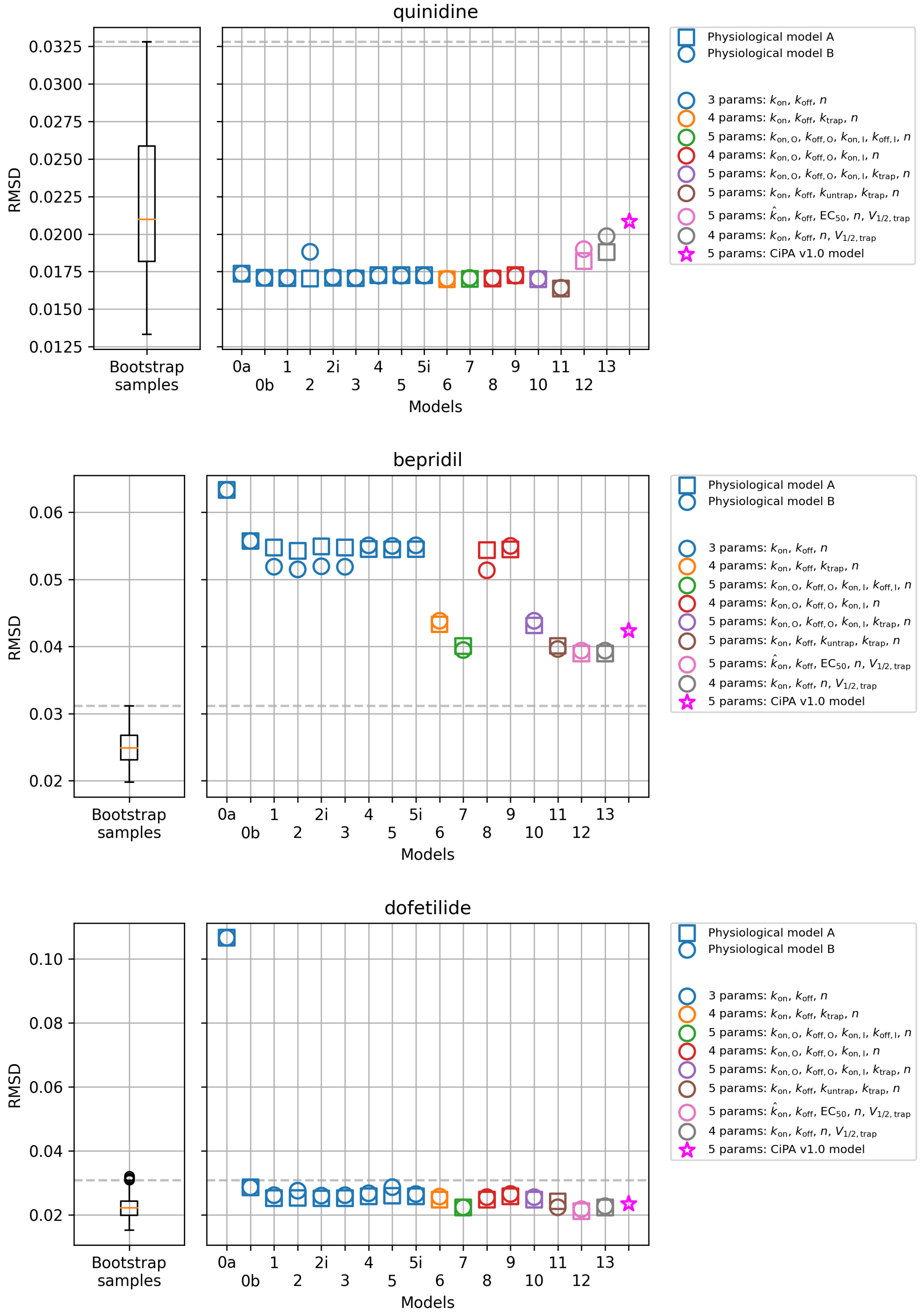


Figure S6: The RMSD of all models compared to the RMSD of the bootstrap samples of the data (box-plot) and the reference binding model (red star, Li et al., 2017). Both base models A (squares) and B (circles) are shown for comparison.

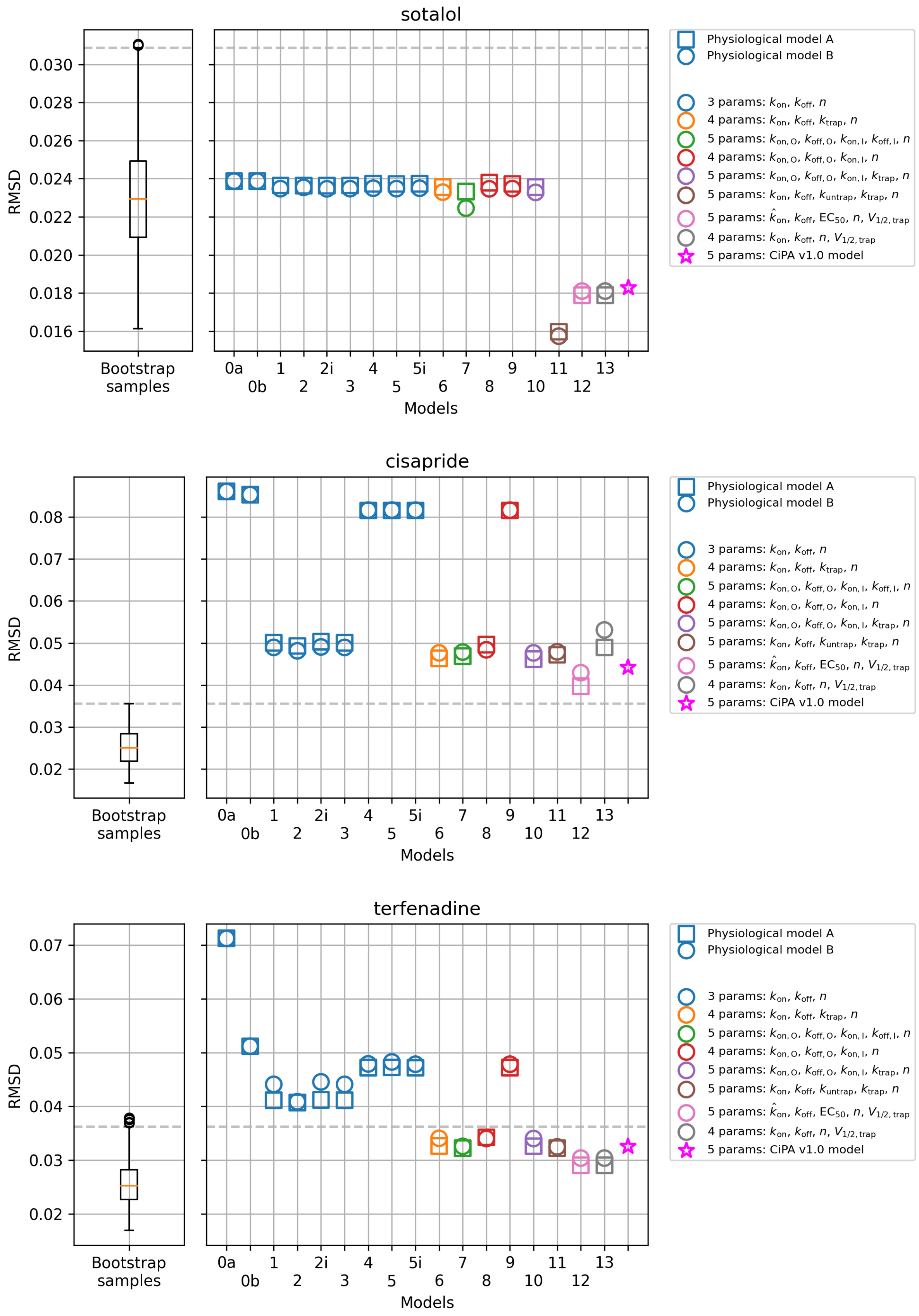


Figure S7: The RMSD of all models compared to the RMSD of the bootstrap samples of the data (box-plot) and the reference binding model (red star, Li et al., 2017). Both base models A (squares) and B (circles) are shown for comparison.

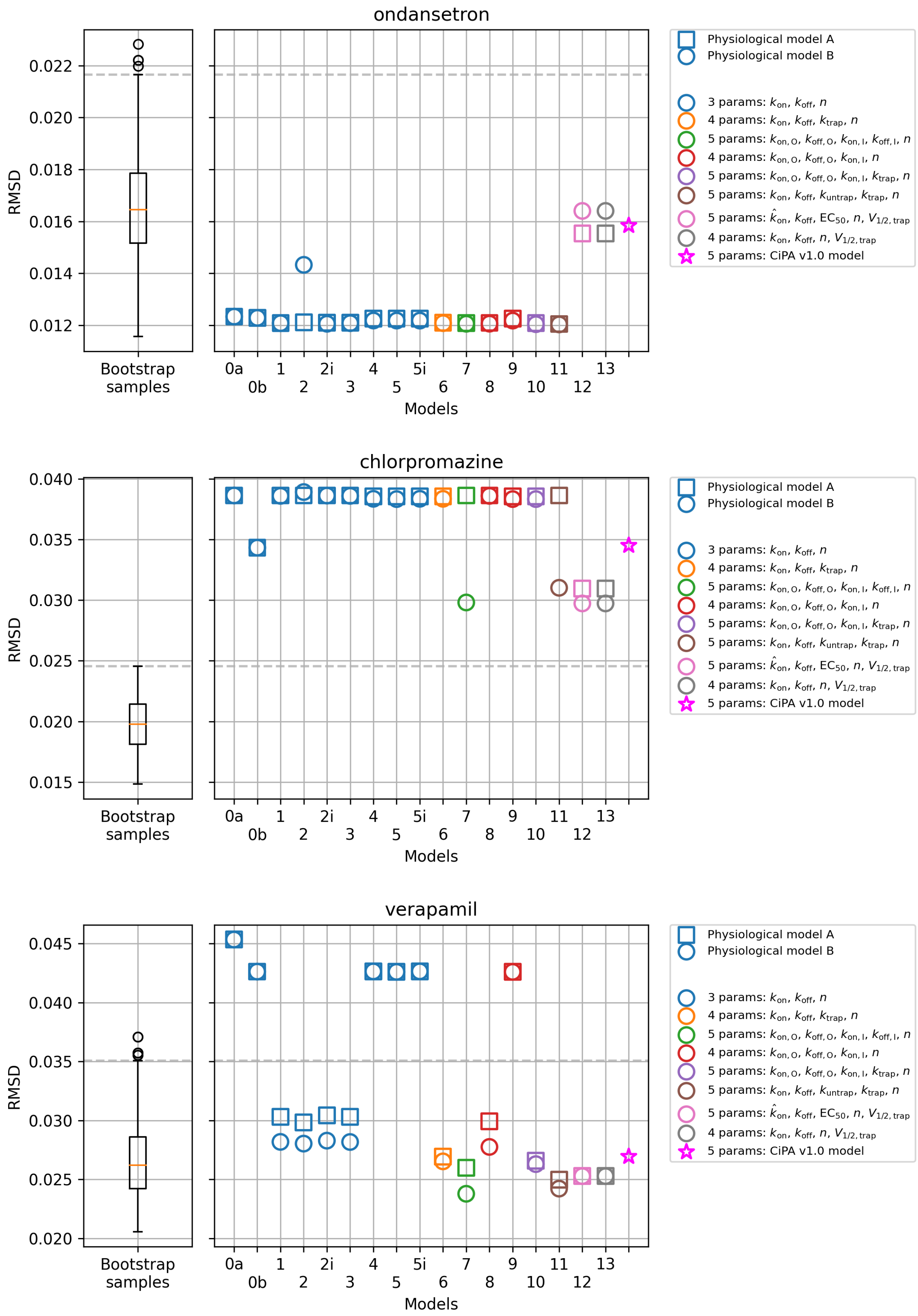


Figure S8: The RMSD of all models compared to the RMSD of the bootstrap samples of the data (box-plot) and the reference binding model (red star, Li et al., 2017). Both base models A (squares) and B (circles) are shown for comparison.

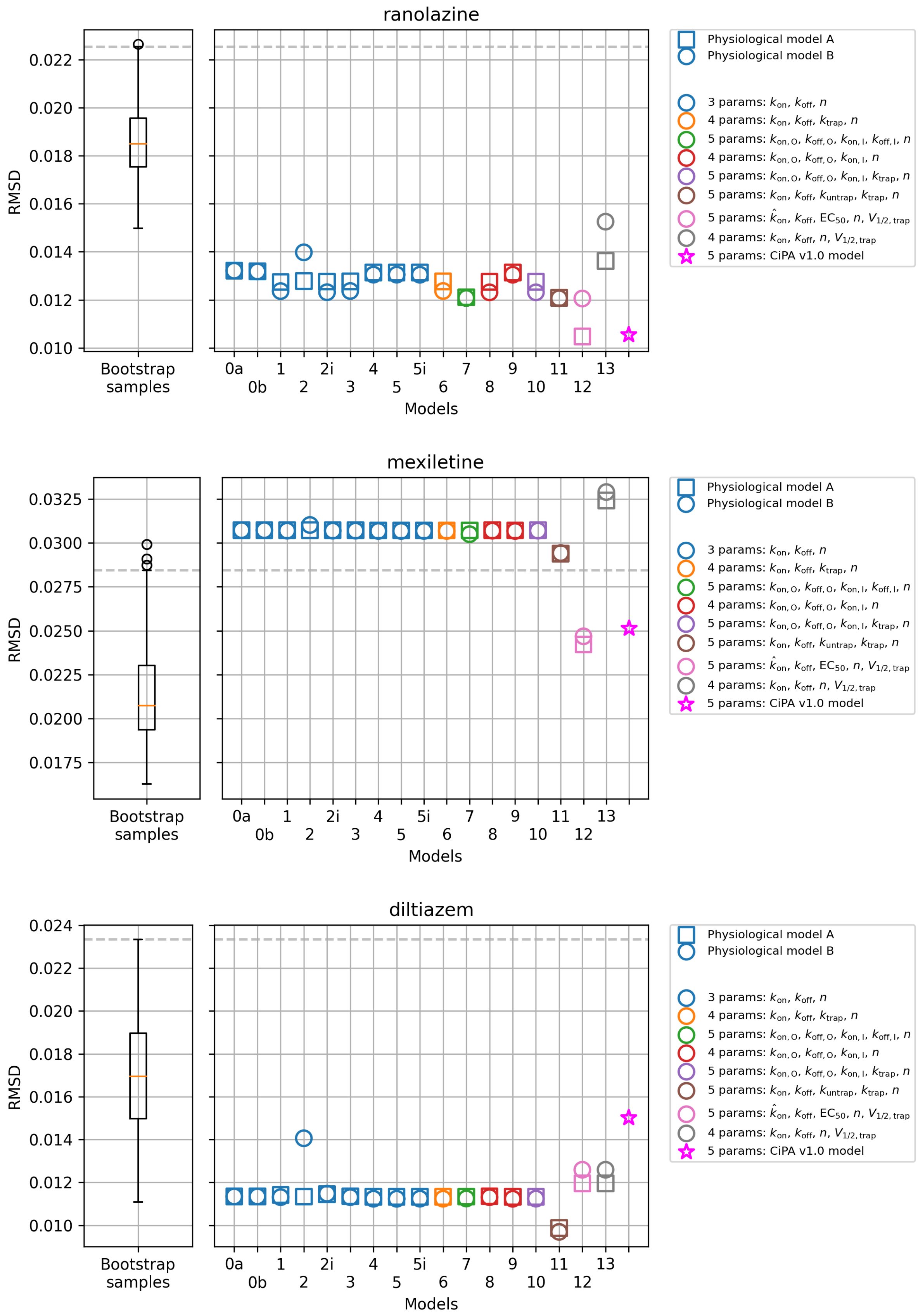


Figure S9: The RMSD of all models compared to the RMSD of the bootstrap samples of the data (box-plot) and the reference binding model (red star, Li et al., 2017). Both base models A (squares) and B (circles) are shown for comparison.

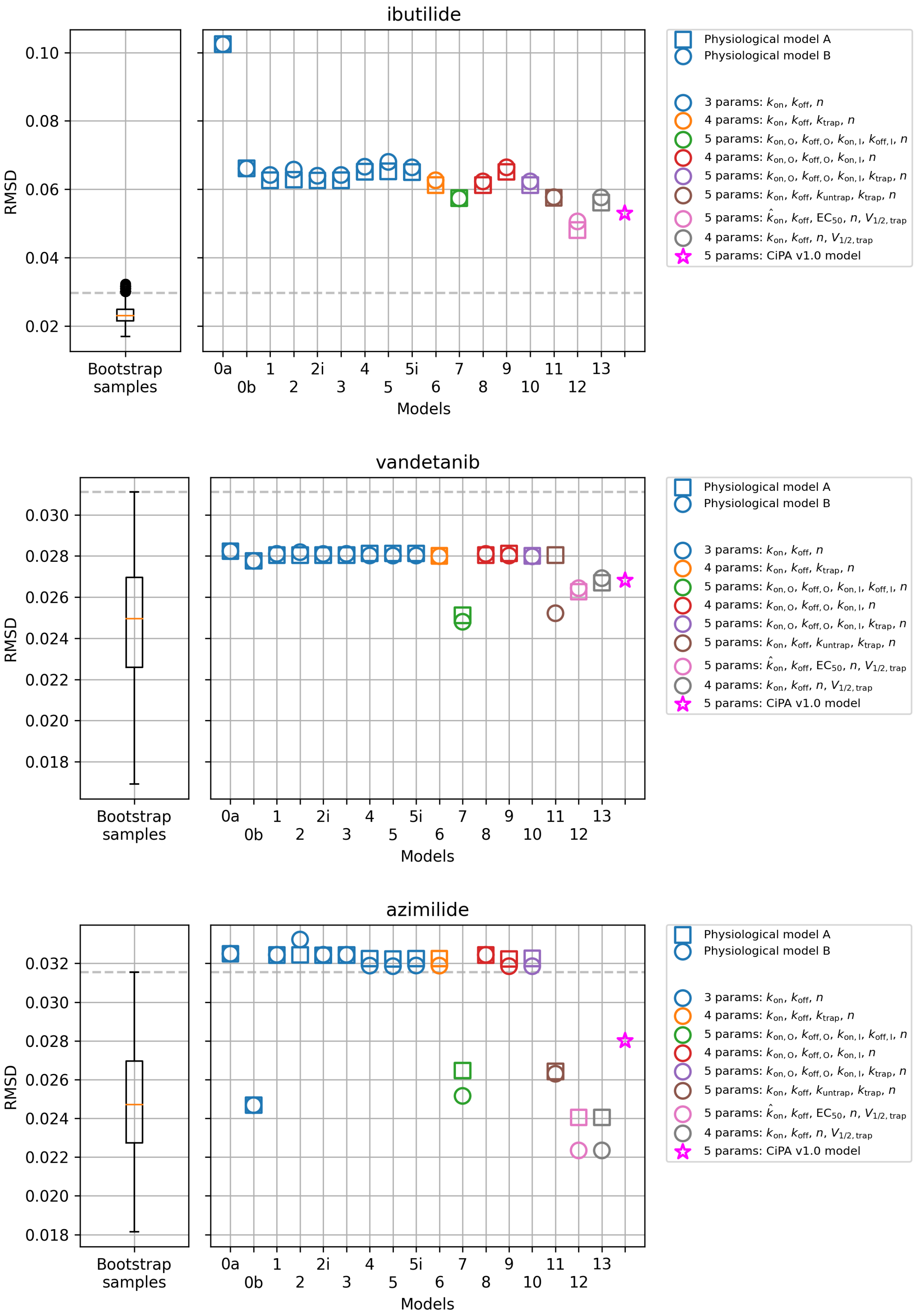


Figure S10: The RMSD of all models compared to the RMSD of the bootstrap samples of the data (box-plot) and the reference binding model (red star, Li et al., 2017). Both base models A (squares) and B (circles) are shown for comparison.

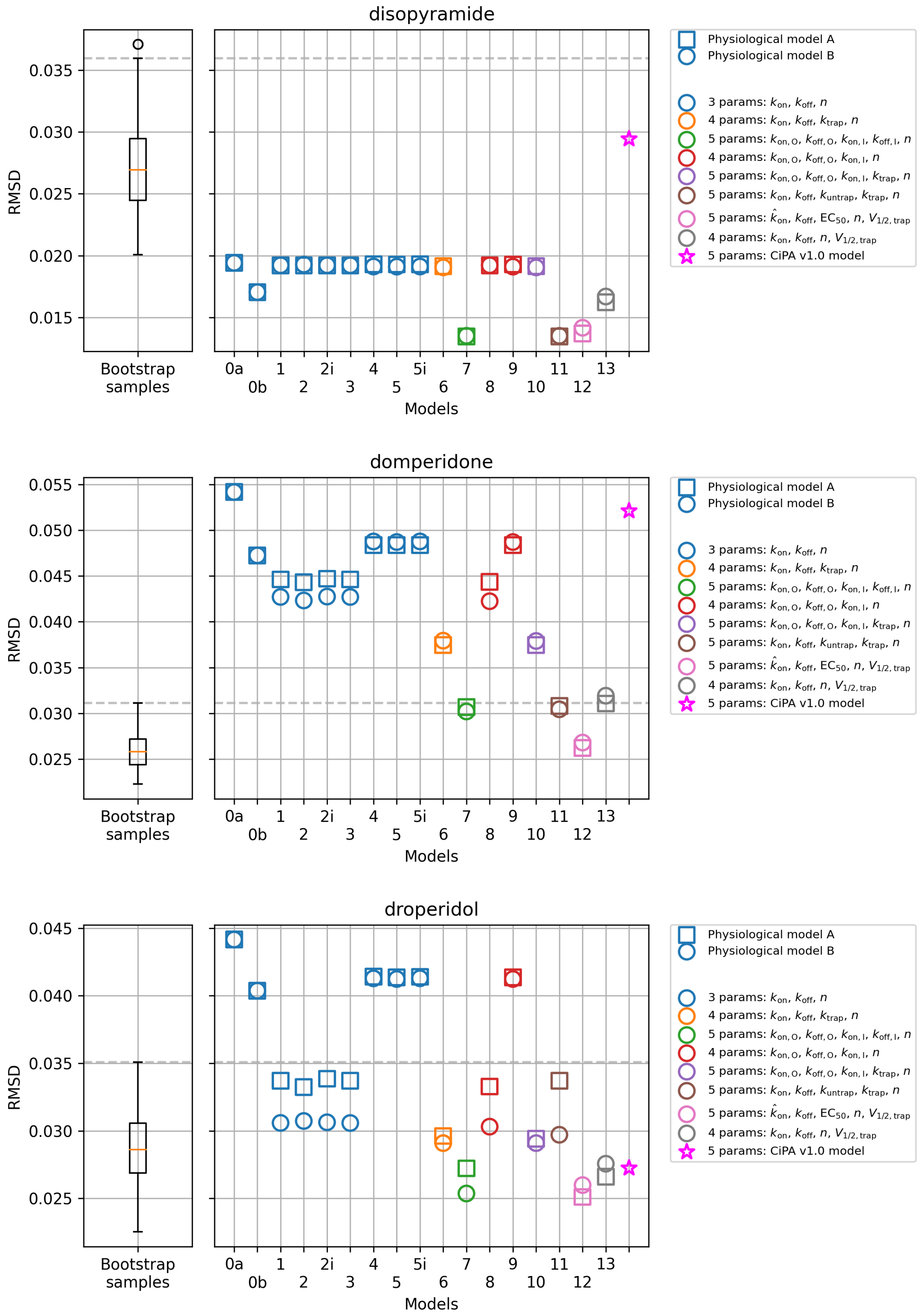


Figure S11: The RMSD of all models compared to the RMSD of the bootstrap samples of the data (box-plot) and the reference binding model (red star, Li et al., 2017). Both base models A (squares) and B (circles) are shown for comparison.

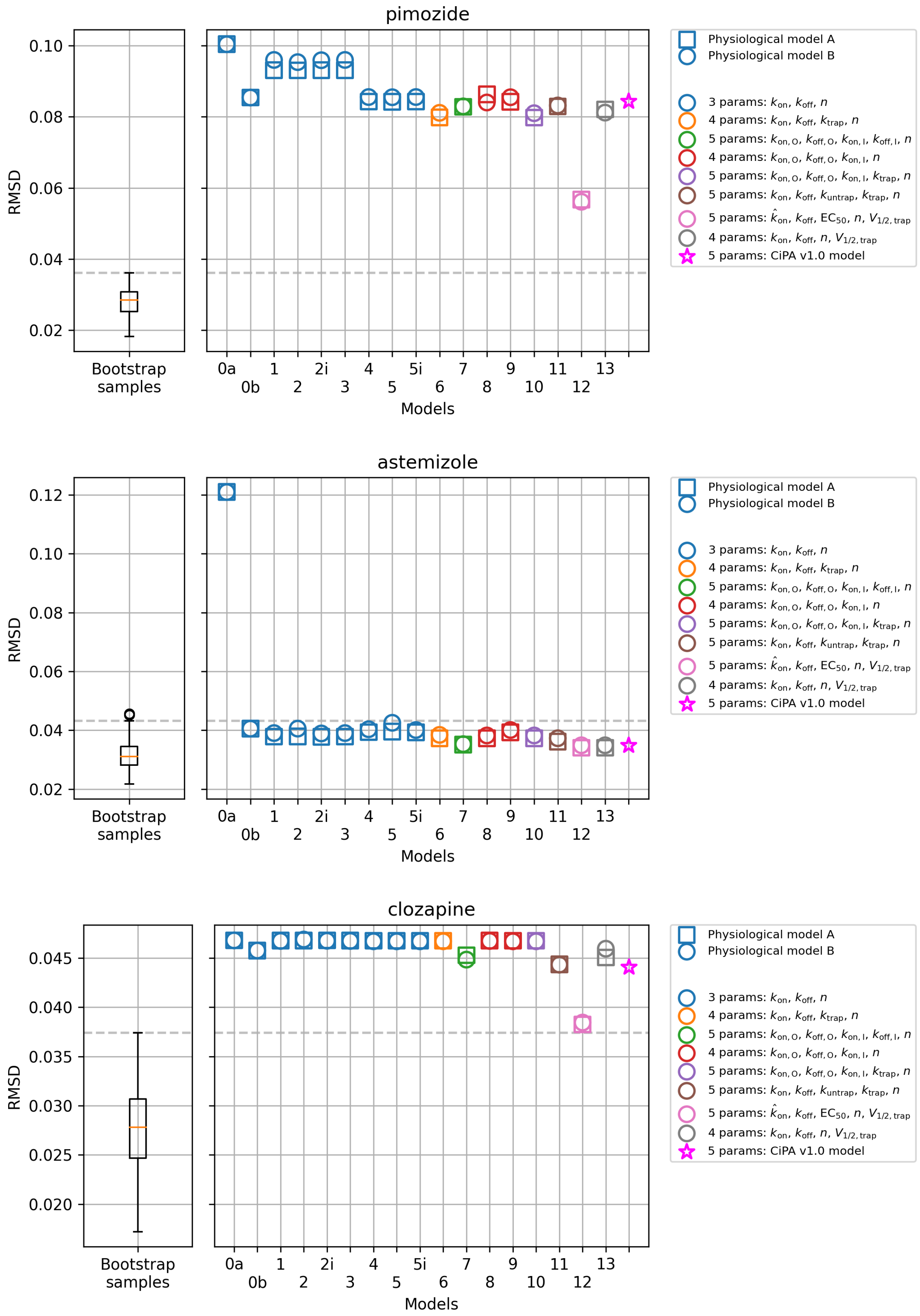


Figure S12: The RMSD of all models compared to the RMSD of the bootstrap samples of the data (box-plot) and the reference binding model (red star, Li et al., 2017). Both base models A (squares) and B (circles) are shown for comparison.

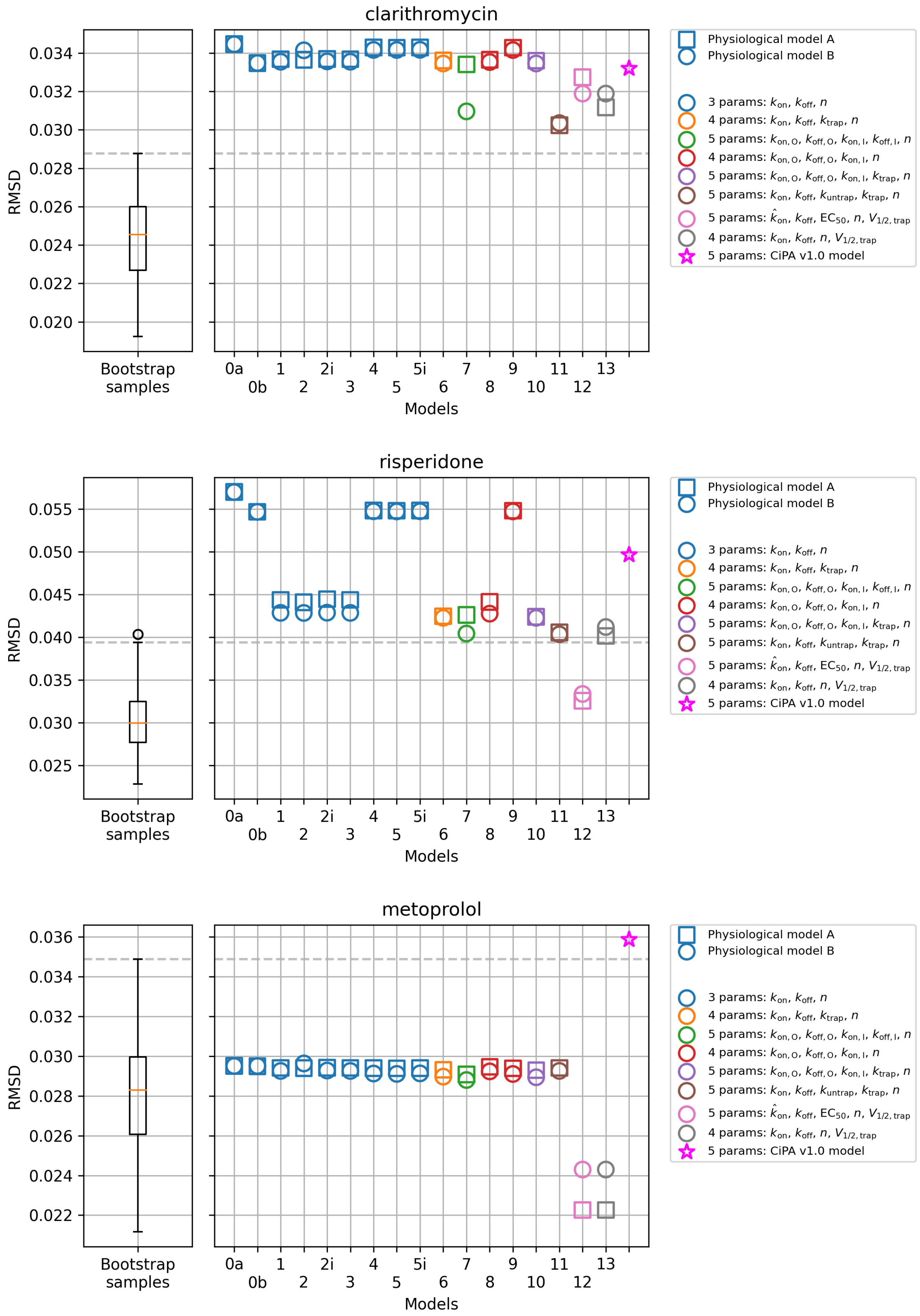


Figure S13: The RMSD of all models compared to the RMSD of the bootstrap samples of the data (box-plot) and the reference binding model (red star, Li et al., 2017). Both base models A (squares) and B (circles) are shown for comparison.

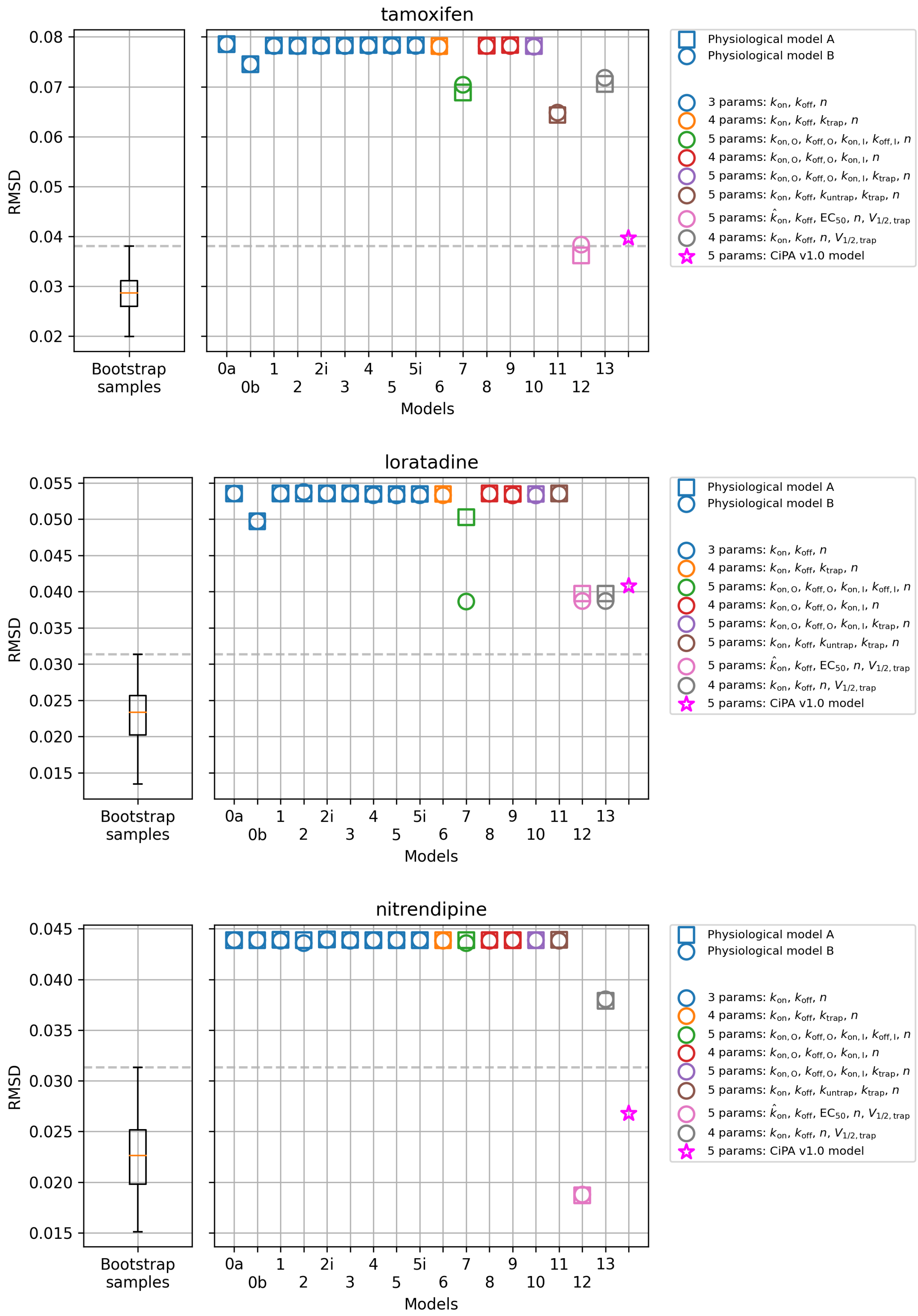


Figure S14: The RMSD of all models compared to the RMSD of the bootstrap samples of the data (box-plot) and the reference binding model (red star, Li et al., 2017). Both base models A (squares) and B (circles) are shown for comparison.

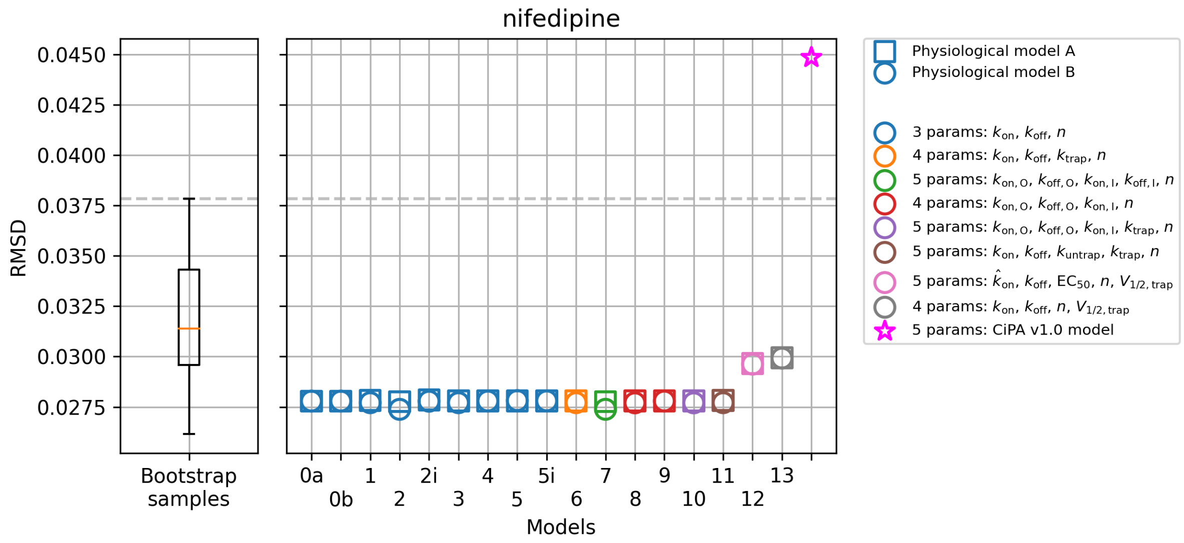


Figure S15: The RMSD of all models compared to the RMSD of the bootstrap samples of the data (box-plot) and the reference binding model (red star, Li et al., 2017). Both base models A (squares) and B (circles) are shown for comparison.

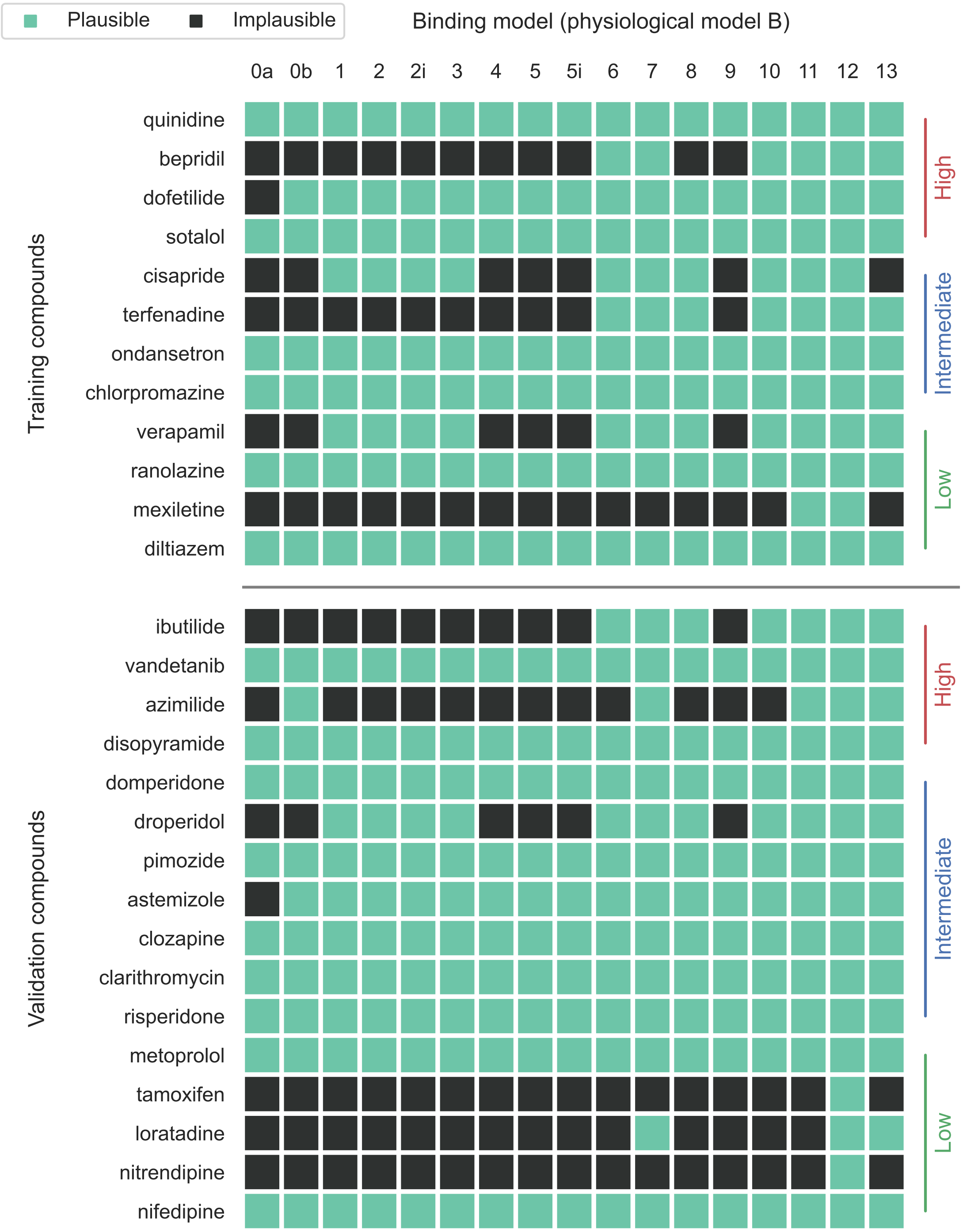


Figure S16: Summary of the selected binding models with physiological model B for all drugs through the RMSD comparison. A binding model (column) is considered to be appropriate for a drug (row)—a plausible model—if coloured in green, where the RMSD of the model to the averaged data is either smaller than the RMSD of the bootstrap samples of data or similar to the CiPA v1.0 model to the averaged data. Model 12 is identical to the drug binding component in the CiPA v1.0 model. Drugs are sorted according to the training and validation lists, and their proarrhythmic risks.

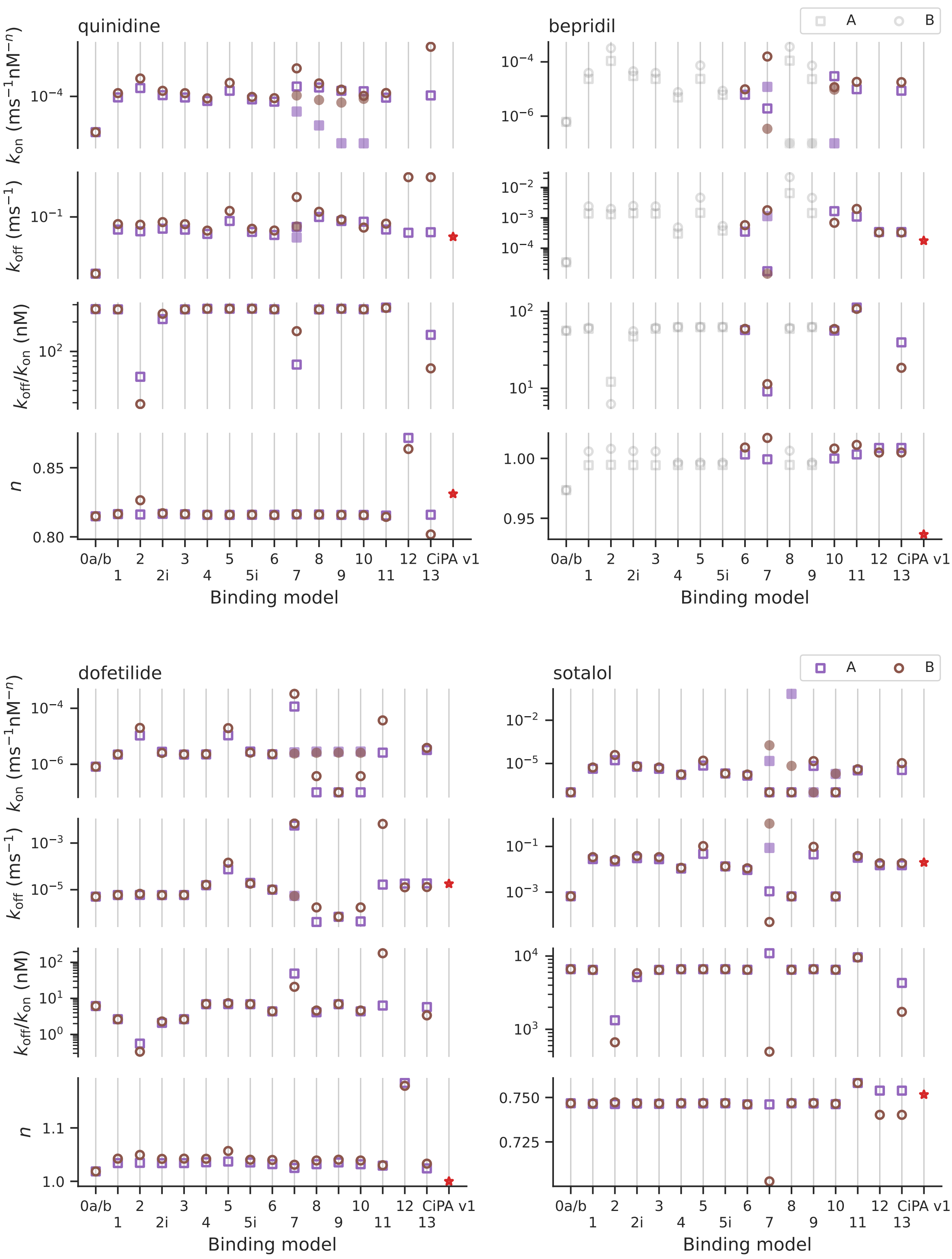


Figure S17: Binding rate parameters $k_{on}$, unbinding rates $k_{off}$, and the Hill coefficients $n$ of the calibrated binding models. Both base models A (purple squares) and B (brown circles) are shown for comparison. Models 7–9 have independent binding and unbinding rates for open and inactivated states; filled squares/circles are the rates for the inactivated states. The models in grey are ruled out through the RMSD comparison. Model 12 is identical to the drug binding component in the reference model (red star).

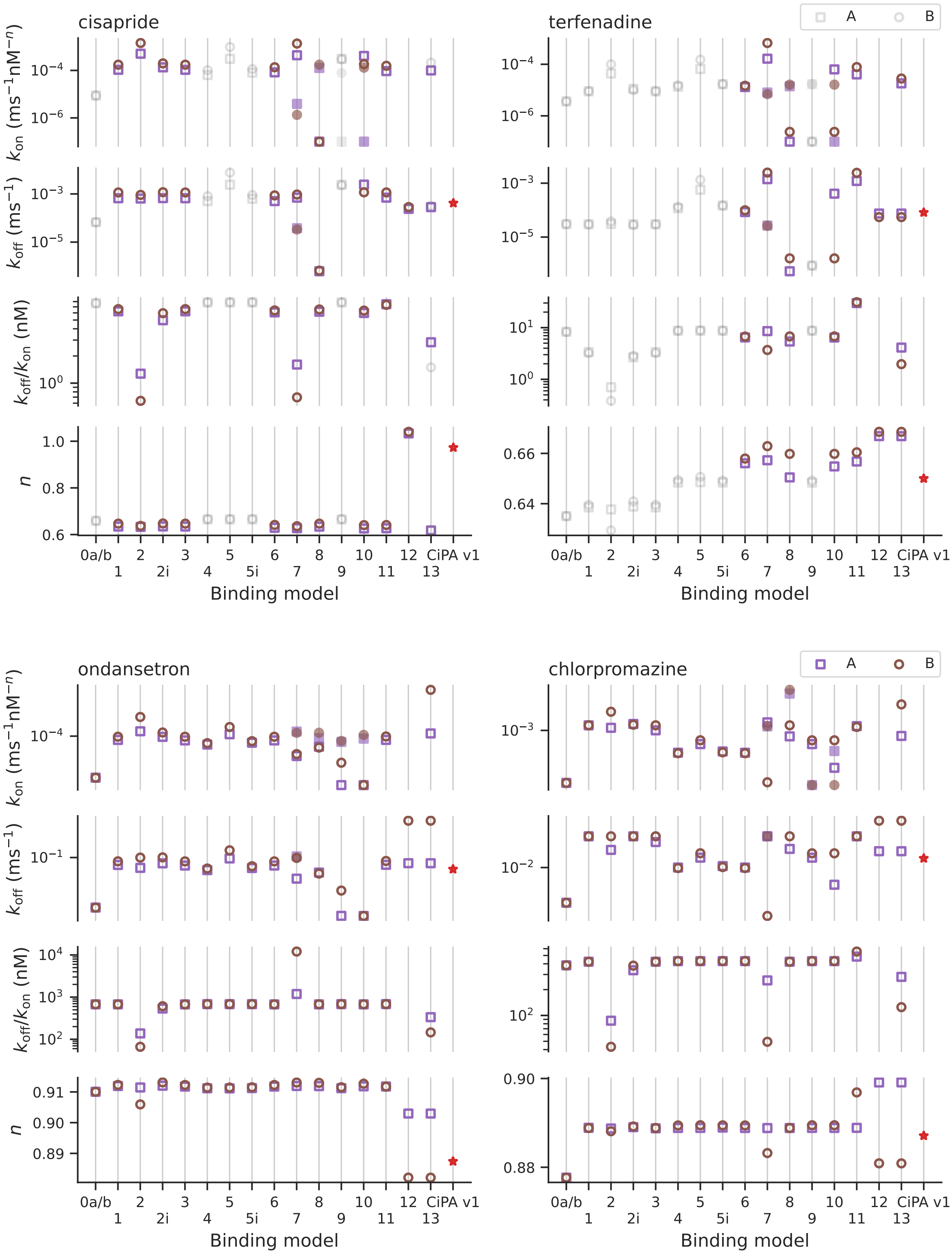


Figure S18: Binding rate parameters $k_{on}$, unbinding rates $k_{off}$, and the Hill coefficients $n$ of the calibrated binding models. Both base models A (purple squares) and B (brown circles) are shown for comparison. Models 7–9 have independent binding and unbinding rates for open and inactivated states; filled squares/circles are the rates for the inactivated states. The models in grey are ruled out through the RMSD comparison. Model 12 is identical to the drug binding component in the reference model (red star).

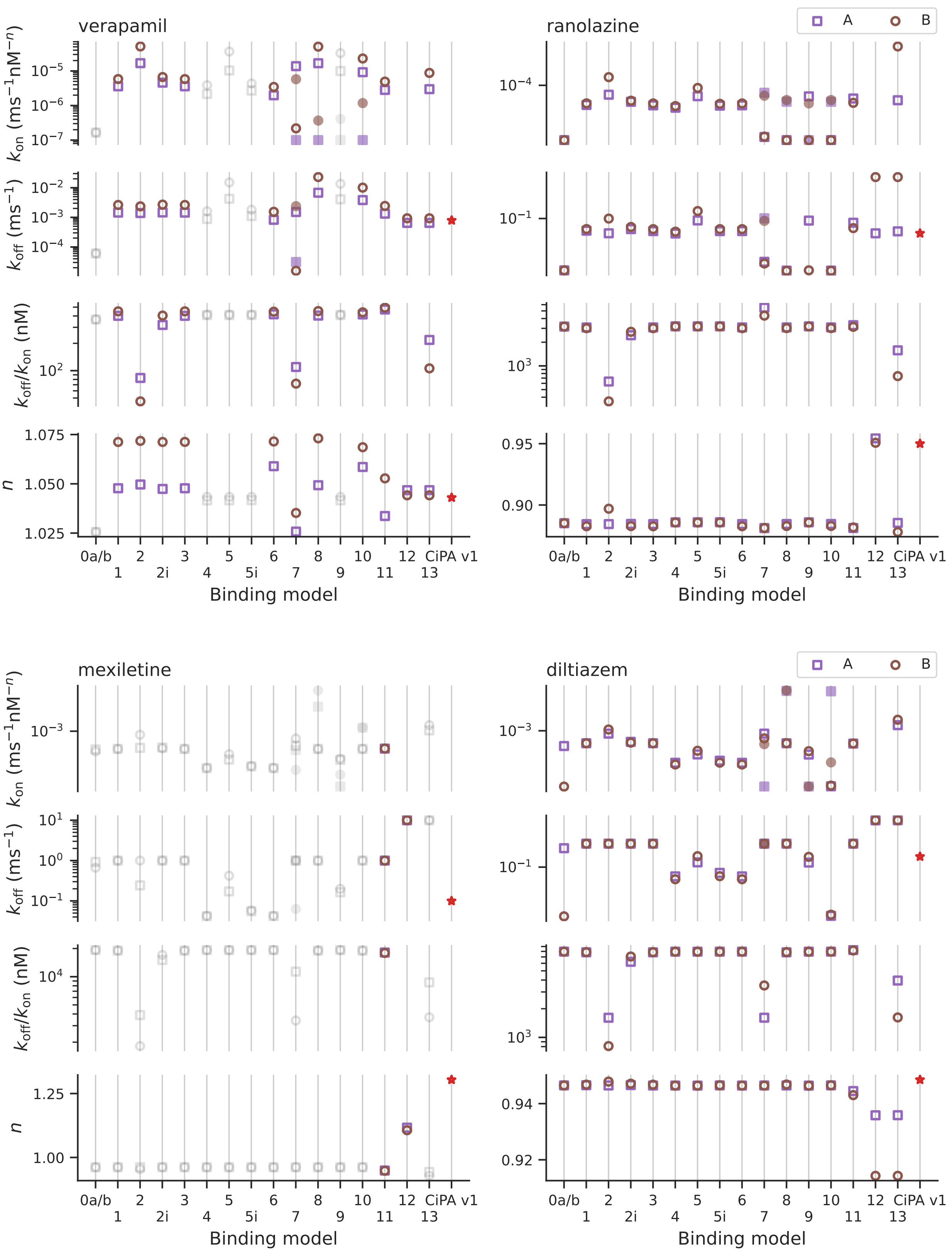


Figure S19: Binding rate parameters $k_{on}$, unbinding rates $k_{off}$, and the Hill coefficients $n$ of the calibrated binding models. Both base models A (purple squares) and B (brown circles) are shown for comparison. Models 7–9 have independent binding and unbinding rates for open and inactivated states; filled squares/circles are the rates for the inactivated states. The models in grey are ruled out through the RMSD comparison. Model 12 is identical to the drug binding component in the reference model (red star).

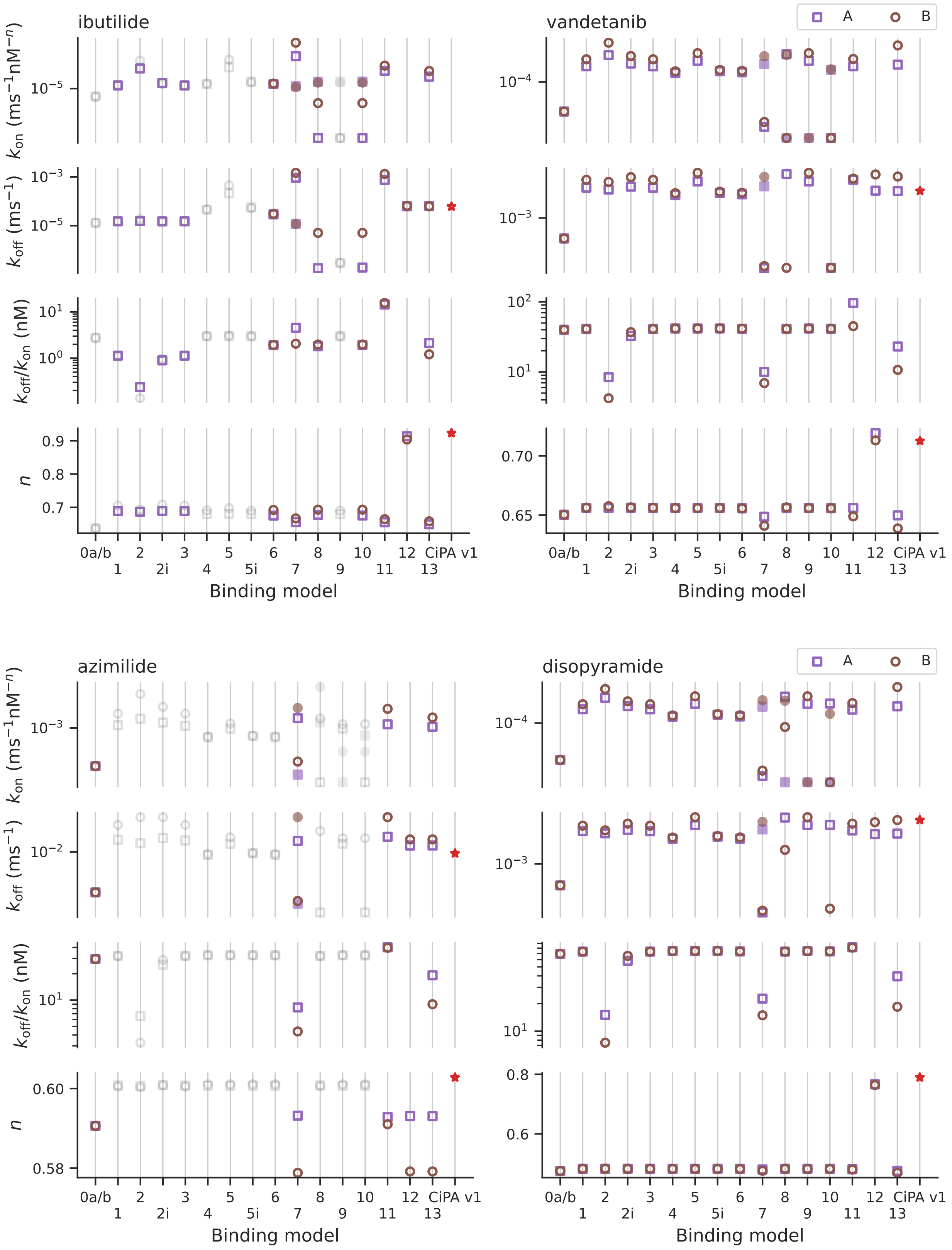


Figure S20: Binding rate parameters $k_{\text{on}}$, unbinding rates $k_{\text{off}}$, and the Hill coefficients $n$ of the calibrated binding models. Both base models A (purple squares) and B (brown circles) are shown for comparison. Models 7–9 have independent binding and unbinding rates for open and inactivated states; filled squares/circles are the rates for the inactivated states. The models in grey are ruled out through the RMSD comparison. Model 12 is identical to the drug binding component in the reference model (red star).

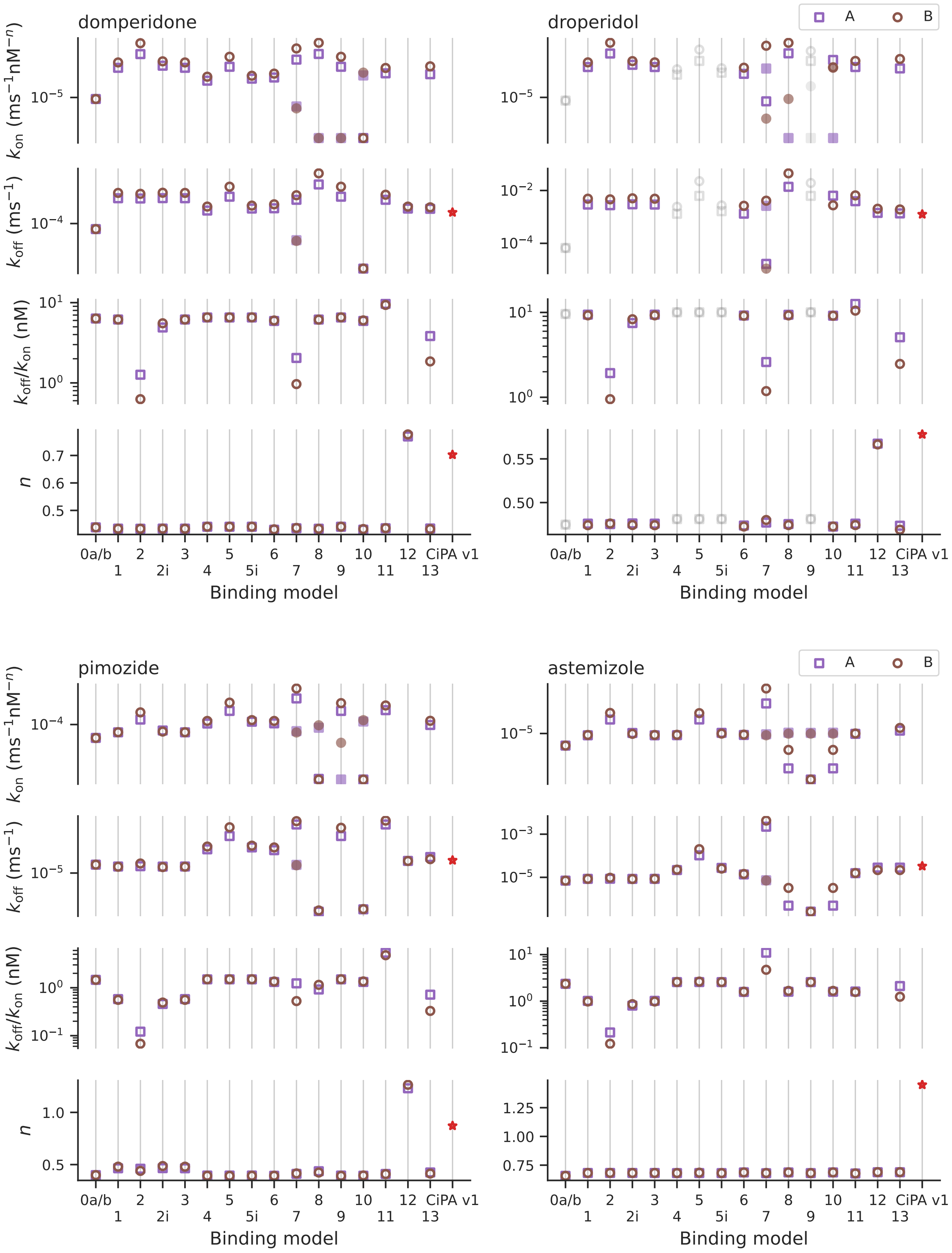


Figure S21: Binding rate parameters $k_{\text{on}}$, unbinding rates $k_{\text{off}}$, and the Hill coefficients $n$ of the calibrated binding models. Both base models A (purple squares) and B (brown circles) are shown for comparison. Models 7–9 have independent binding and unbinding rates for open and inactivated states; filled squares/circles are the rates for the inactivated states. The models in grey are ruled out through the RMSD comparison. Model 12 is identical to the drug binding component in the reference model (red star).

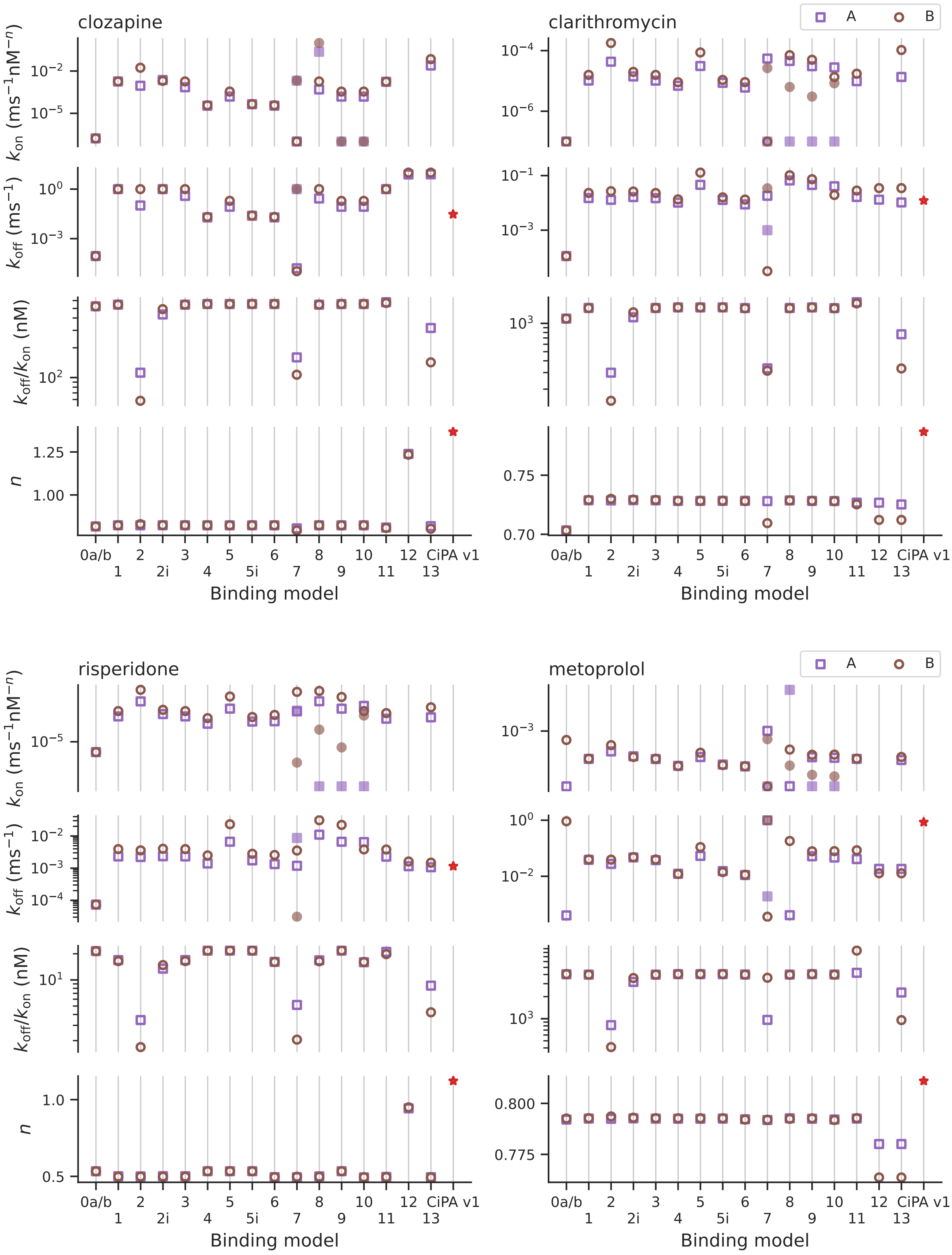


Figure S22: Binding rate parameters $k_{on}$, unbinding rates $k_{off}$, and the Hill coefficients $n$ of the calibrated binding models. Both base models A (purple squares) and B (brown circles) are shown for comparison. Models 7–9 have independent binding and unbinding rates for open and inactivated states; filled squares/circles are the rates for the inactivated states. The models in grey are ruled out through the RMSD comparison. Model 12 is identical to the drug binding component in the reference model (red star).

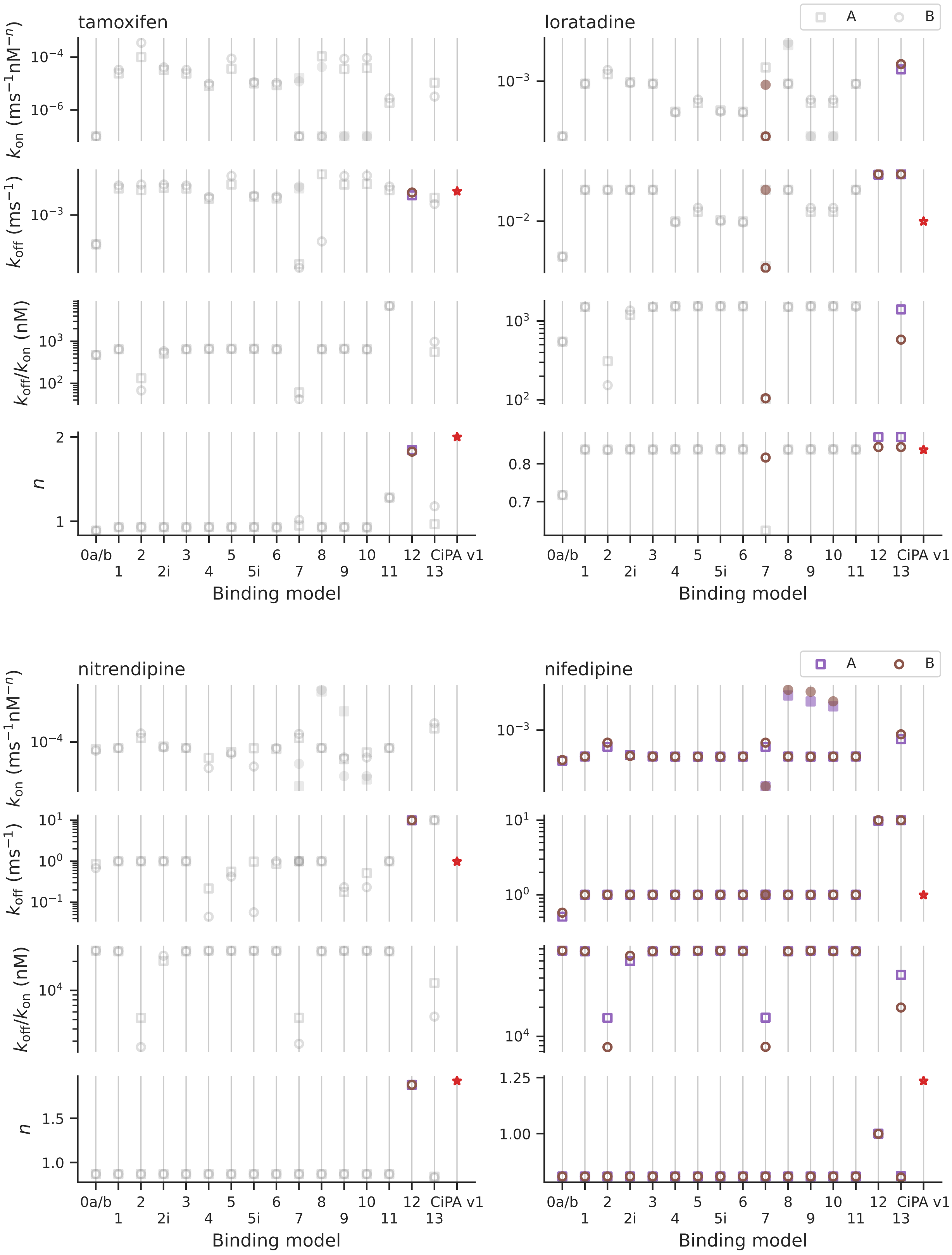


Figure S23: Binding rate parameters $k_{on}$, unbinding rates $k_{off}$, and the Hill coefficients $n$ of the calibrated binding models. Both base models A (purple squares) and B (brown circles) are shown for comparison. Models 7–9 have independent binding and unbinding rates for open and inactivated states; filled squares/circles are the rates for the inactivated states. The models in grey are ruled out through the RMSD comparison. Model 12 is identical to the drug binding component in the reference model (red star).

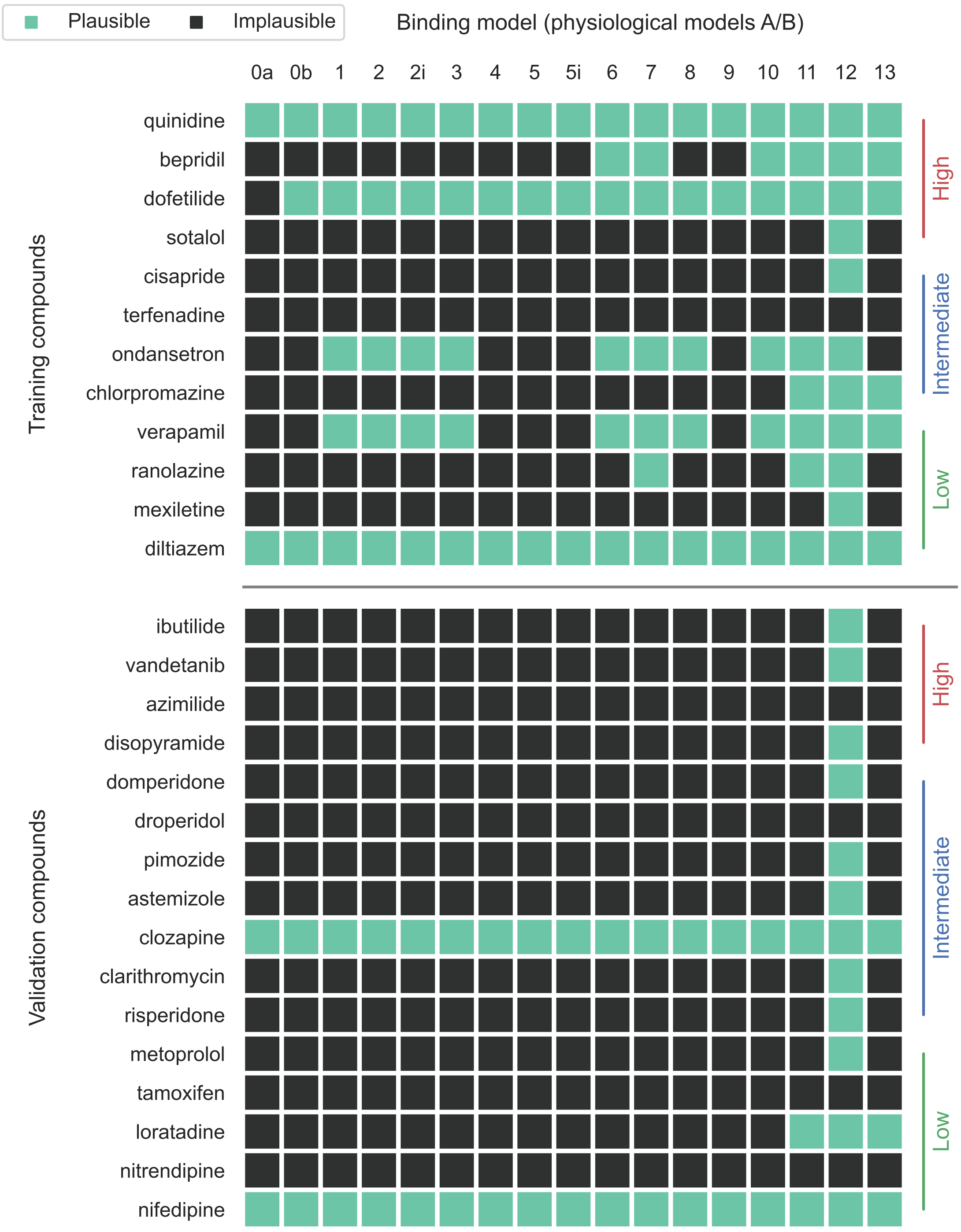


Figure S24: Summary of the selected binding models with physiological models A and B for all drugs through the RMSD comparison, assuming the Hill coefficient $n = 1$. A binding model (column) is considered to be appropriate for a drug (row)—a plausible model—if coloured in green, where the RMSD of the model to the averaged data is either smaller than the RMSD of the bootstrap samples of data or similar to the CiPA v1.0 model to the averaged data. Model 12 is identical to the drug binding component in the CiPA v1.0 model but with $n = 1$. Drugs are sorted according to the training and validation lists, and their proarrhythmic risks.
